# Versatile, Multivalent Nanobody Cocktails Efficiently Neutralize SARS-CoV-2

**DOI:** 10.1101/2020.08.24.264333

**Authors:** Yufei Xiang, Sham Nambulli, Zhengyun Xiao, Heng Liu, Zhe Sang, W. Paul Duprex, Dina Schneidman-Duhovny, Cheng Zhang, Yi Shi

## Abstract

The outbreak of COVID-19 has severely impacted global health and the economy. Cost-effective, highly efficacious therapeutics are urgently needed. Here, we used camelid immunization and proteomics to identify a large repertoire of highly potent neutralizing nanobodies (Nbs) to the SARS-CoV-2 spike (S) protein receptor-binding domain (RBD). We discovered multiple elite Nbs with picomolar to femtomolar affinities that inhibit viral infection at sub-ng/ml concentration, more potent than some of the best human neutralizing antibodies. We determined a crystal structure of such an elite neutralizing Nb in complex with RBD. Structural proteomics and integrative modeling revealed multiple distinct and non-overlapping epitopes and indicated an array of potential neutralization mechanisms. Structural characterization facilitated the bioengineering of novel multivalent Nb constructs into multi-epitope cocktails that achieved ultrahigh neutralization potency (IC50s as low as 0.058 ng/ml) and may prevent mutational escape. These thermostable Nbs can be rapidly produced in bulk from microbes and resist lyophilization, and aerosolization. These promising agents are readily translated into efficient, cost-effective, and convenient therapeutics to help end this once-in-a-century health crisis.

## Introduction

Globally a novel, highly transmissible coronavirus severe acute respiratory syndrome coronavirus 2 (SARS-CoV-2) {Zhu, 2020 #1; Zhou, 2020 #2} has infected more than 20 million people and has claimed over 700,000 lives, with the numbers still rising as of Aug 2020. Despite preventive measures, such as quarantines and lock-downs that help curb viral transmission, the virus rebounds after lifting social restrictions. Safe and effective therapeutics and vaccines remain in dire need.

Like other zoonotic coronaviruses, SARS-CoV-2 expresses a surface spike (S) glycoprotein, which consists of S1 and S2 subunits forming a homotrimeric viral spike to interact with host cells. The interaction is mediated by the S1 receptor-binding domain (RBD), which binds the peptidase domain (PD) of angiotensin-converting enzyme-2 (hACE2) as a host receptor {Wrapp, 2020 #3}. Structural studies have revealed different conformations of the spike {Walls, 2020 #4; Cai, 2020 #5}. In the pre-fusion stage, the RBD switches between an inactive, closed conformation, and an active open conformation necessary for interacting with hACE2. In the post-fusion stage, S1 dissociates from the trimer, while S2 undergoes a dramatic conformational change to trigger host membrane fusion {Fan, 2020 #28}. Most recently, investigations into COVID-19 convalescence individuals’ sera have led to the identification of highly potent neutralizing IgG antibodies (NAbs) primarily targeting the RBD but also non-RBD epitopes {Cao, 2020 #6; Robbiani, 2020 #7; Hansen, 2020 #8; Liu, 2020 #9; Brouwer, 2020 #10; Chi, 2020 #19; Wec, 2020 #65}. High-quality NAbs may overcome the risks of the Fc-associated antibody-dependent enhancement (ADE) and are promising therapeutic and prophylactic candidates {Zohar, 2020 #11; Eroshenko, 2020 #12}.

The V_H_H antibodies or nanobodies (Nbs) are minimal, monomeric antigen-binding domains derived from camelid single-chain antibodies {Muyldermans, 2013 #13}. Unlike IgG antibodies, Nbs are small (~15 kDa), highly soluble and stable, readily bioengineered into bi/multivalent forms, and are amenable to low-cost, efficient microbial production. Due to their robust physicochemical properties, Nbs can be administered by inhalation, making their use against the respiratory viruses very appealing {Vanlandschoot, 2011 #14; Detalle, 2016 #15}. Previous efforts have yielded broadly neutralizing Nbs for various challenging viruses, including Dengue, RSV, and HIV-1 {Vanlandschoot, 2011 #15; Weiss, 2019 #29; Rossey, 2017 #66}. Very recently, several SARS-CoV-2 neutralizing Nbs have been identified, by screening SARS-CoV or Middle East respiratory syndrome (MERS) cross-reacting Nbs or using synthetic Nb libraries for RBD binding. However, these synthetic Nbs generally neutralize the virus at μg to sub-μg/ml concentration {Huo, 2020 #16; Wrapp, 2020 #17; Konwarh, 2020 #18;Custódio, 2020 #35;Walter, 2020 #36;Gai, 2020 #3;Gai, 2020 #38}, which are hundreds of times less potent than the best NAbs, likely due to monovalency and lack of affinity maturation {Barnes, 2020 #13}. The development of high-quality anti-SARS-CoV-2 Nbs may provide a novel means for versatile, cost-effective therapeutics and point-of-care diagnosis.

We have developed and characterized a collection of diverse, soluble, stable, and high-affinity camelid Nbs that target the RBD. The majority of the high-affinity RBD Nbs efficiently neutralize SARS-CoV-2. A subset has exceptional neutralization potency, comparable with, or better than the most potent human NAbs. We used integrative structural approach and mapped multiple epitopes of neutralizing Nbs. We then determined an atomic structure of an elite Nb of sub-ng/ml neutralization potency in complex with the RBD. Our results revealed that while highly potent neutralizing Nbs predominantly recognize the concave, hACE2 binding site, efficient neutralization can also be accomplished through other RBD epitopes. Finally, structural characterizations facilitated the bioengineering of multivalent Nbs into multi-epitope cocktails that achieved remarkable neutralization potencies of as low as 0.058 ng/ml (1.3 pM), which may be sufficient to prevent the generation of escape mutants.

## Results

### Development of highly potent SARS-CoV-2 neutralizing Nbs

To produce high-quality SARS-CoV-2 neutralizing Nbs, we immunized a llama with the recombinant RBD protein expressed in human 293T cells. Compared to the pre-bleed, after affinity maturation, the post-immunized serum showed potent and specific serologic activities towards RBD binding with a titer of 1.75×10^6^ (**Fig S1a**). The serum efficiently neutralized the pseudotyped SARS-CoV-2 at the half-maximal neutralization titer (NT50) of 310,000 (**Fig S1b**), orders of magnitude higher than the convalescent sera obtained from recovered COVID-19 patients {Cao, 2020 #6; Robbiani, 2020 #7}. To further characterize these activities, we separated the single-chain V_H_H antibodies from the IgG antibodies from the serum. We confirmed that the single-chain antibodies achieve specific, high-affinity binding to the RBD and possess sub-nM half-maximal inhibitory concentration (IC50 = 509 pM) against the pseudotyped virus (**Fig S1c**).

We identified thousands of high-affinity V_H_H Nbs from the RBD-immunized llama serum using a robust proteomic strategy that we have recently developed {Xiang, 2020 #67} (**Fig S2a**). This repertoire includes ~350 unique CDR3s (complementarity-determining regions). For *E.coli* expression, we selected 109 highly diverse Nb sequences from the repertoire with unique CDR3s to cover various biophysical, structural, and potentially different antiviral properties of our Nb repertoire. Ninety four Nbs were purified and tested for RBD binding by ELISA, from which we confirmed 71 RBD-specific binders (**Fig S2b-c, Table S1**). Of these RBD-specific binders, 49 Nbs presented high solubility and high-affinity (ELISA IC50 below 30 nM, **Fig 1a**), and were promising candidates for functional characterizations. We used a SARS-CoV-2-GFP pseudovirus neutralization assay to screen and characterize the antiviral activities of these high-affinity Nbs. Interestingly, the vast majority (94%) of the tested Nbs can neutralize the pseudotype virus below 3 μM (**Fig 1b**). 90% of them blocked the pseudovirus below 500 nM. Only 20-40% of high-affinity RBD-specific mAbs identified from the patients’ sera have been reported to possess comparable potency {Robbiani, 2020 #7; Cao, 2020 #6}. Over three quarters (76%) of the Nbs efficiently neutralized the pseudovirus below 50 nM, and 6% had neutralization activities below 0.5 nM. Finally we tested the potential of 14 to neutralize SARS-CoV-2 Munich strain using our recently published PRNT50 assay {Klimstra, 2020 #73}. All the Nbs reached 100% neutralization and neutralized the virus in a dose-dependent manner. The IC50s span from single-digit ng/ml to sub-ng/ml, with three unusual neutralizers of Nbs 89, 20, and 21 to be 2.1 ng/ml (0.137 nM), 1.6 ng/ml (0.102 nM), and 0.7 ng/ml (0.045 nM), respectively, based on the pseudovirus assay (**Fig 1c, 1e**). Similar values (0.154 nM, 0.048 nM, and 0.021 nM, for Nbs 89, 20, and 21) were reproducibly obtained using SARS-CoV-2 (**Fig 1d, 1e**). There was an excellent correlation between the two neutralization assays (R^2^= 0.92, **Fig S3**).

**Figure 1.**
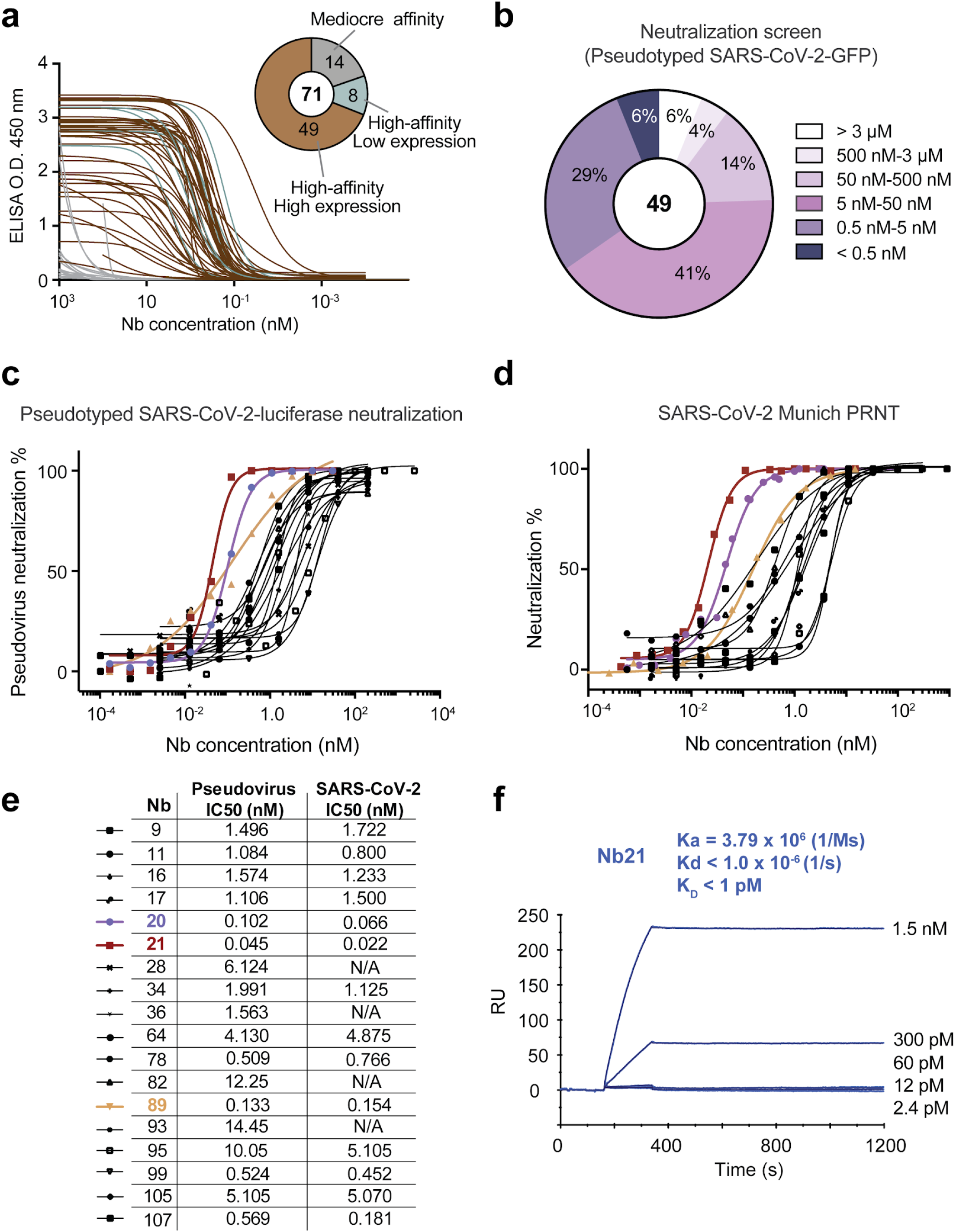
Production and characterizations of high-affinity RBD Nbs for SARS-CoV-2 neutralization. **1a**: The binding affinities of 71 Nbs towards RBD by ELISA. The pie chart shows the number of Nbs according to affinity and solubility. **1b**: Screening of 49 soluble, high-affinity Nbs by SARS-CoV-2-GFP pseudovirus neutralization assay. n = 1 for Nbs with neutralization potency IC50 <= 50 nM, n = 2 for Nbs with neutralization potency IC50 > 50 nM. **1c**: The neutralization potency of 18 highly potent Nbs was calculated based on the pseudotyped SARS-CoV-2 neutralization assay (luciferase). Purple, red, and yellow lines denote Nbs 20, 21, and 89 with IC50 < 0.2 nM. Two different purifications of the pseudovirus were used. The average neutralization percentage was shown for each data point (n = 5 for Nbs 20, 21; n=2 for all other Nbs). **1d**: The neutralization potency of the top 14 neutralizing Nbs by SARS-CoV-2 plaque reduction neutralization test (PRNT). The average neutralization percentage was shown for each data point (n=4 for Nbs 20, 21, and 89; n=2 for other Nbs). **1e:** A table summary of pseudotyped and SARS-CoV-2 neutralization potency for 18 Nbs. N/A: not tested. **1f**: The SPR binding kinetics measurement of Nb21.

We measured the binding kinetics of Nbs 89, 20, and 21 by surface plasmon resonance (SPR) (**Fig S4a-b**). While Nbs 89 and 20 have an affinity of 108 pM and 10.4 pM, our best-neutralizing Nb21 did not show detectable dissociation from the RBD during 20 min SPR analysis. The femtomolar affinity of Nb21 potentially explains its unusual neutralization potency (**Fig 1f**). We determined the thermostability of the top three neutralizing Nbs (89, 20, and 21) from the *E.coli* periplasmic preparations to be 65.9, 71.8, and 72.8°C, respectively (**Fig S4c**). Finally, we tested the on-shelf stability of Nb21, which remained soluble after ~ 6 weeks of storage at room temperature after purification and could well tolerate lyophilization. No multimeric forms or aggregations were detected by size-exclusion chromatography (SEC) (**Fig S4d**). Together these results suggest that these neutralizing Nbs have the necessary physicochemical properties required for advanced therapeutic applications.

### Integrative structural characterization of the Nb neutralization epitopes

Epitope mapping based on atomic resolution structure determination by X-ray crystallography and Cryo-Electron Microscopy (CryoEM) is highly accurate but low-throughput. Here, we integrated information from SEC, cross-linking mass spectrometry (CXMS), shape and physicochemical complementarity, and statistics to determine structural models of RBD-Nb complexes {Rout, 2019 #20; Yu, 2017 #39; Leitner, 2016 #40; Chait, 2016 #41}. First, we performed SEC experiments to distinguish between Nbs that share the same epitope as Nb21 (thus complete with Nb 21 on an SEC) and those that bind to non-overlapping epitopes. Nbs 9, 16, 17, 20, 64, 82, 89, 99 and 107 competed with Nb21 for RBD binding based on SEC profiles (**Fig 2a, S5**), indicating that their epitopes significantly overlap. In contrast, higher mass species (from early elution volumes) corresponding to the trimeric complexes composed of Nb21, RBD, and one of the Nbs (34, 36, 93, 105, and 95) were evident (**Fig 2b, S6a-h**). Moreover, Nb105 competed with Nb34 and Nb95, which did not compete for RBD interaction, suggesting the presence of two distinct and non-overlapping epitopes. Second, we cross-linked Nb-RBD complexes by DSS (disuccinimidyl suberate) and identified on average, four intermolecular cross-links by MS for Nbs 20, 93, 34, 95, and 105. The cross-links were used to map the RBD epitopes derived from the SEC data (**Methods**). Our cross-linking models identified five epitopes (I, II, III, IV, and V corresponding to Nbs 20, 93, 34, 95, and 105) (**Fig 2c**). The models satisfied 90% of the cross-links with an average precision of 7.8 Å (**Fig 2d**, **Table S2**). Our analysis confirmed the presence of a dominant Epitope I (e.g., epitopes of Nbs 20 and 21) overlapping with the hACE2 binding site. Epitope II also co-localized with the hACE2 binding site, while epitopes III-V did not (**Fig 2e**). Interestingly, epitope I Nbs had significantly shorter CDR3 (four amino acids shorter, p = 0.005) than other epitope binders (**Fig S6i**). Despite this, the vast majority of the selected Nbs potently inhibited the virus with an IC50 below 30 ng/ml (2 nM) (**Table S1**).

**Figure 2.**
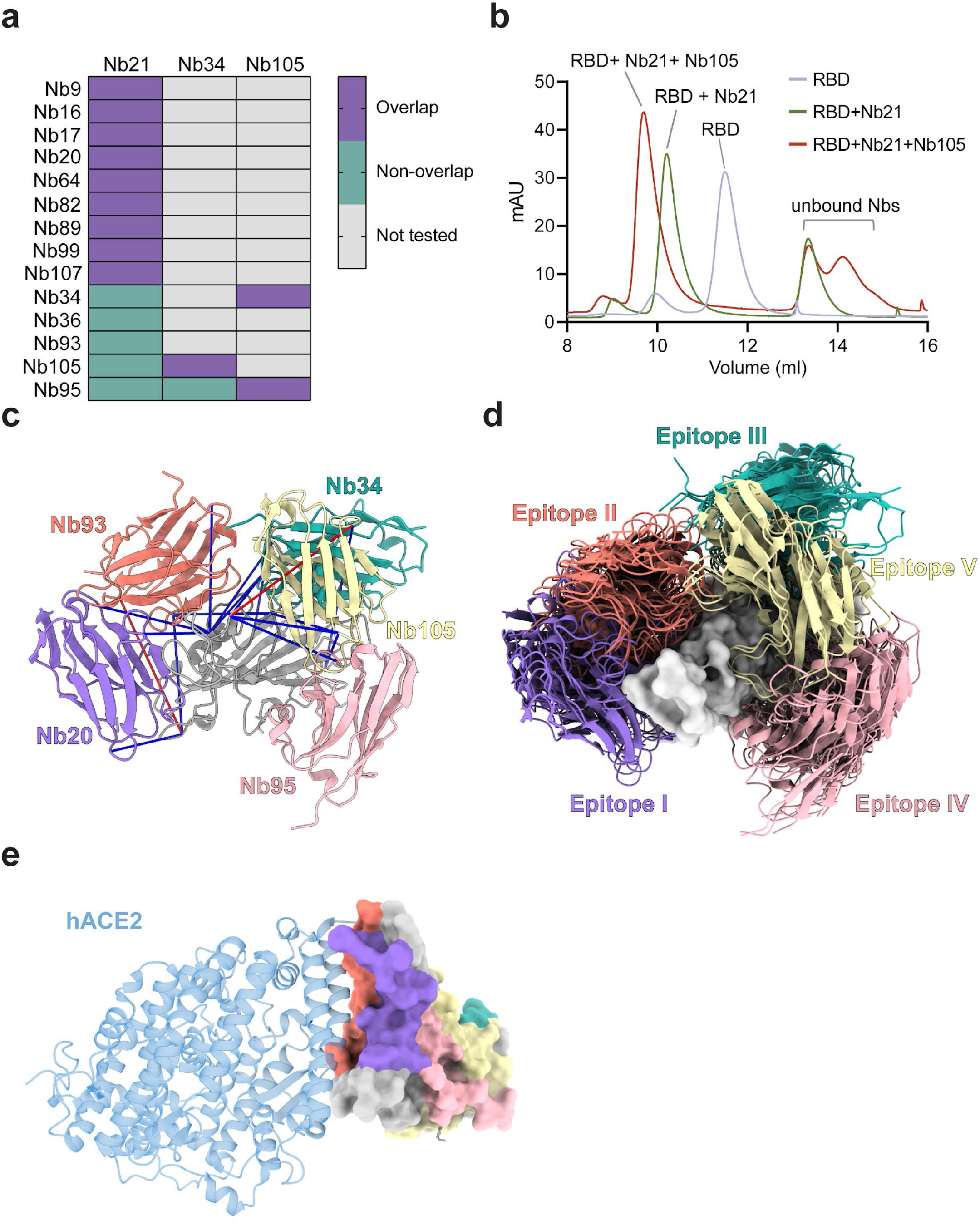
Nb epitope mapping by integrative structural proteomics. **Fig 2a:** A summary of Nb epitopes based on size exclusion chromatography (SEC) analysis. Light salmon color: Nbs that bind the same RBD epitope. Sea green: Nbs of different epitopes. **Fig 2b:** A representation of SEC profiling of RBD, RBD-Nb21 complex, and RBD-Nb21-Nb105 complex. The y-axis represents UV 280 nm absorbance units (mAu). **Fig 2c:** A cartoon model showing the localization of five Nbs that bind different epitopes: Nb20 (medium purple), Nb34 (light sea green), Nb93 (salmon), Nb105 (pale goldenrod) and Nb95 (light pink) in complex with the RBD (gray). Blue and red lines represent DSS cross-links shorter or longer than 28Å, respectively. **Fig 2d:** Top 10 scoring cross-linking based models for each Nb (cartoons) on top of the RBD surface. **Fig 2e**: The surface display of different Nb neutralization epitopes on the RBD in complex with hACE2 (cartoon model in blue).

### Crystal structure of RBD-Nb20 and analysis of Nbs 20 and 21 interactions with RBD

To explore the molecular mechanisms that underlie the unusually potent neutralization activities of Epitope I Nbs, we determined a crystal structure of the RBD-Nb20 complex at a resolution of 3.3 Å by molecular replacement (**Methods, Table S3**). Most of the residues in RBD (N334-G526) and the entire Nb20, particularly those at the protein interaction interface, are well resolved. There are two copies of RBD-Nb20 complexes in one asymmetric unit, which are almost identical with an RMSD of 0.277 Ȧ over 287 Cα atoms. In the structure, all three CDRs of Nb20 interact with the RBD by binding to its large extended external loop with two short β-strands (**Fig 3a**) {Wang, 2020 #23}. E484 of RBD forms hydrogen bonding and ionic interactions with the side chains of R31 (CDR1) and Y104 (CDR3) of Nb20, while Q493 of RBD forms hydrogen bonds with the main chain carbonyl of A29 (CDR1) and the side chain of R97 (CDR3) of Nb20. These interactions constitute a major polar interaction network at the RBD and Nb20 interface. R31 of Nb20 also engages in a cation-π interaction with the side chain of F490 of the RBD (**Fig 3b**). In addition, M55 from the CDR2 of Nb20 packs against residues L452, F490, and L492 of RBD to form hydrophobic interactions at the interface). Another small patch of hydrophobic interactions is formed among residues V483 of RBD and F45 and L59 from the framework β-sheet of Nb20 (**Fig 3c**).

**Figure 3.**
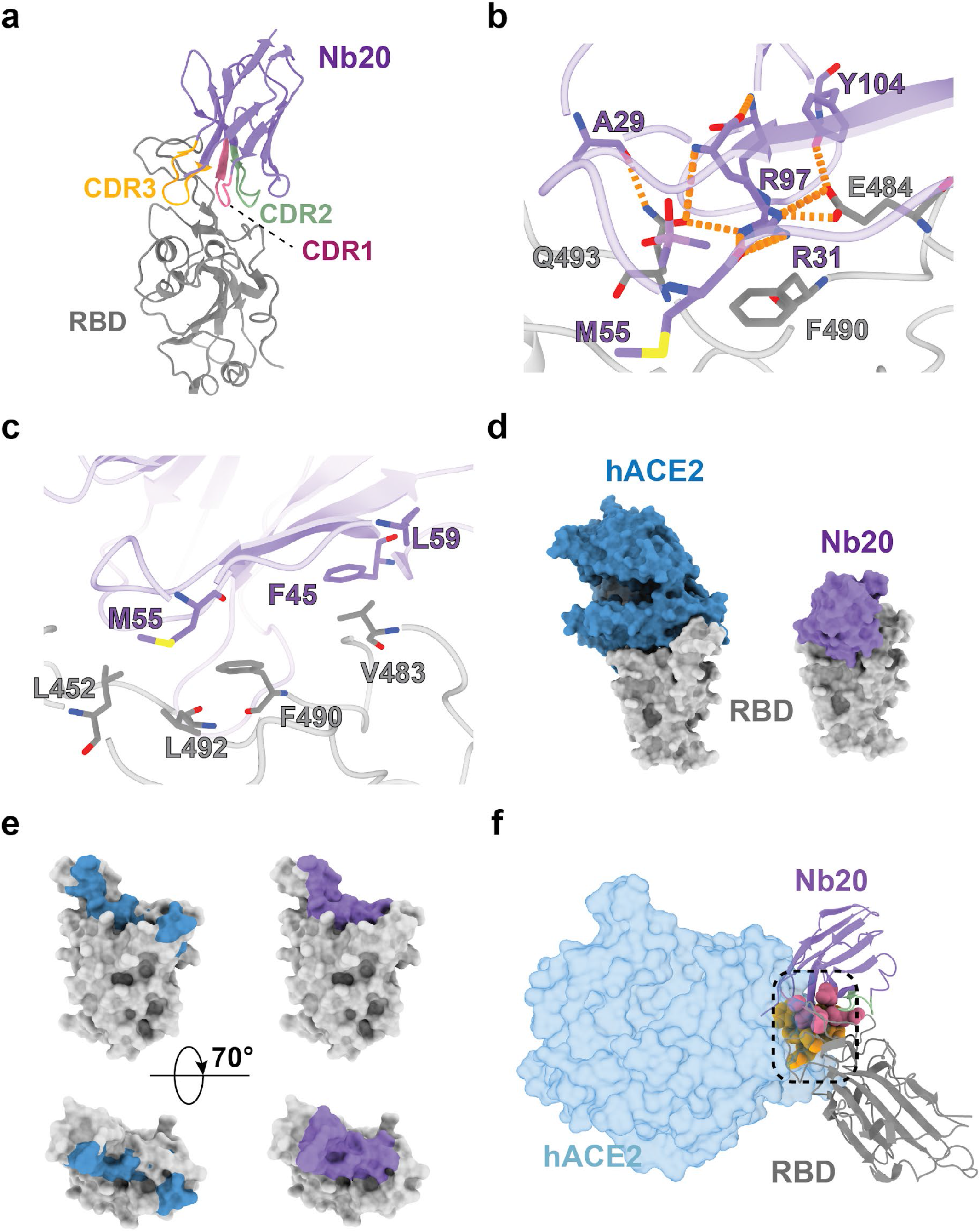
Crystal structure analysis of an ultrahigh affinity Nb in complex with the RBD. **3a.** Cartoon presentation of Nb20 in complex with the RBD. CDR1, 2, and 3 are in red, green, and orange, respectively. **3b.** Zoomed-in view of an extensive polar interaction network that centers around R35 of Nb20. **3c.** Zoomed-in view of hydrophobic interactions. **3d.** Surface presentation of the Nb20-RBD and hACE2-RBD complex (PDB: 6M0J) **3e.** Surface presentation of RBD with hACE2 binding epitope colored in steel blue and Nb20 epitope colored in medium purple. **3f.** The CDR1 and CDR3 residues (medium violet pink and light goldenrod in spheres, respectively) of Nb20 overlap with hACE2 binding site (light blue) on the RBD (gray).

The binding mode of Nb20 to the RBD is distinct from all other reported SARS-CoV-2 neutralizing Nbs, which generally recognize similar epitopes in the RBD external loop region {Li, 2020 #33; Walter, 2020 #34; Huo, 2020 #18} (**Fig S7**). The extensive hydrophobic and polar interactions (**Fig 3b-c**) between the RBD and Nb20 stem from the remarkable shape complementarity (**Fig 3d**) between all the CDRs and the external RBD loop, leading to ultrahigh-affinity (~ 10 pM). We further modeled the structure of the best neutralizer Nb21 with RBD based on our crystal structure (**Methods**). Only four residues vary between Nb20 and Nb21 (**Fig S8a**), all of which are on CDRs. Two substitutions are at the RBD binding interface. S52 and M55 in the CDR2 of Nb20 are replaced by two asparagine residues N52 and N55 in Nb21. In our modeled structure, N52 forms a new H-bond with N450 of RBD (**Fig S8b**). While N55 does not engage in additional interactions with RBD, it creates a salt bridge with the side chain of R31, which stabilizes the polar interaction network among R31 and Y104 of Nb21 and Q484 of RBD (**Fig S8b**). All of those may contribute to a slower off-rate of Nb21 vs. Nb20 (**Fig 2f, S4a**) and stronger neutralization potency. Structural comparison of RBD-Nb20/21 and RBD-hACE2 (PDB 6LZG) {Wang, 2020 #23} clearly showed that the interfaces for Nb20/21 and hACE2 partially overlap (**Fig 3d-e**). Notably, the CDR1 and CDR3 of Nb20/21 would severely clash with the first helix of hACE2, the primary binding site for RBD (**Fig 3f**). Our high-resolution structural study provides novel insight into the exceptional binding affinities of epitope I Nbs that are likely to contribute to the sub-ng/ml neutralization capability.

### Potential mechanisms of SARS-CoV-2 neutralization by Nbs

To understand the outstanding antiviral efficacy of our Nbs better, we superimposed RBD-Nb complexes to different spike conformations based on cryoEM structures. We found that three copies of Nb20/21 can simultaneously bind all three RBDs in their “down” conformations (PDB 6VXX) {Walls, 2020 #4} that correspond to the inactive spike (**Fig 4b**). Our analysis indicates a potential mechanism by which Nbs 20 and 21 (Epitope I) lock RBDs in their down conformation with ultrahigh affinity. Combined with the steric interference with hACE2 binding in the RBD open conformation (**Fig 4a**), these mechanisms may explain the exceptional neutralization potencies of Epitope I Nbs.

**Figure 4.**
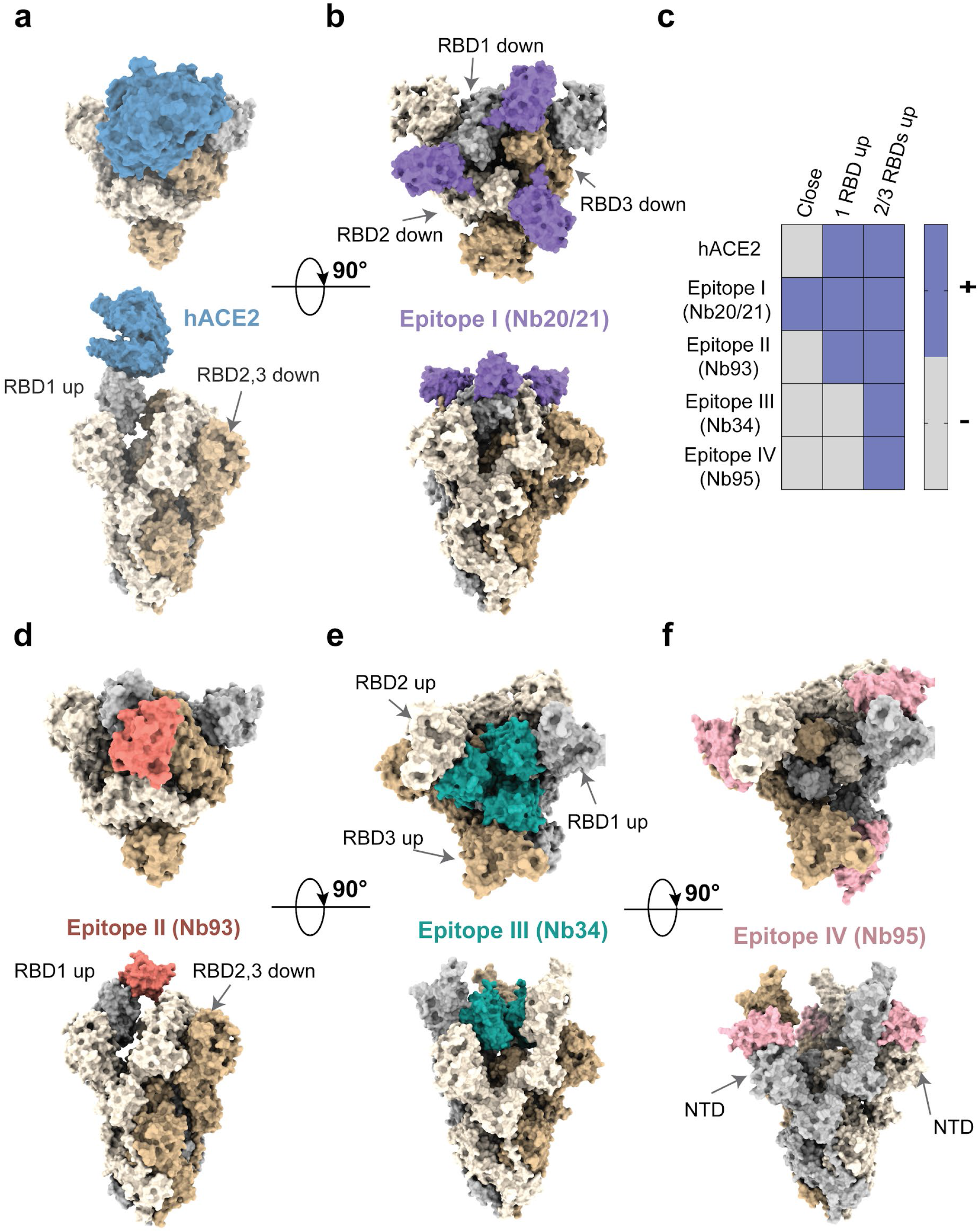
Potential mechanisms of SARS-CoV-2 neutralization by Nbs. **4a.** hACE2 (grey) binding to spike trimer conformation (wheat, plum, and light blue colors) with one RBD up (PDBs 6VSB, 6LZG). **4b.** Nb20 (Epitope I, medium purple) partially overlaps with the hACE2 binding site and can bind the closed spike conformation with all RBDs down (PDB 6VXX). **4c.** A summary of spike conformations accessible (+) to the Nbs of different epitopes. **4d.** Nb93 (Epitope II, salmon) partially overlaps with the hACE2 binding site and can bind to spike conformations with at least one RBD up (PDB 6VSB). **4e-f.** Nb34 (Epitope III, light sea blue) and Nb95 (Epitope IV, light pink) do not overlap with the hACE2 binding site and bind to spike conformations with at least two open RBDs (PDB 6XCN).

Other epitope-binders do not fit into this inactive conformation without steric clashes and appear to utilize different neutralization strategies (**Fig 4c**). For example, Epitope II: Nb 93 co-localizes with hACE2 binding site and can bind the spike in the one RBD “up” conformation (**Fig 4d**, PDB 6VSB) {Wrapp, 2020 #3}. It may neutralize the virus by blocking the hACE2 binding site. Epitope III and IV Nbs can only bind when two or three RBDs are at their “up” conformations (PDB 6XCN) {Barnes, 2020 #13} where the epitopes are exposed. In the all RBDs “up” conformation, three copies of Nbs can directly interact with the trimeric spike. Interestingly, through RBD binding, Epitope III: Nb34 can be accommodated on top of the trimer to lock the helices of S2 in the prefusion stage, preventing their large conformational changes for membrane fusion (**Fig 4e**). When superimposed onto the all “up” conformation, Epitope IV: Nb95 is proximal to the rigid NTD of the trimer, presumably restricting the flexibility of the spike domains (**Fig 4f**).

### Development of flexible, multivalent Nbs for highly efficient viral neutralization

Epitope mapping enabled us to bioengineer a series of multivalent Nbs (**Fig 5a**). Specifically, we designed two sets of constructs that build upon our most potent Nbs. The homotrimeric Nbs, in which a flexible linker sequence (either 31 or 25 amino acids, **Methods**) separates each monomer Nb (such as Nb21 or Nb20), were designed to increase the antiviral activities through avidity binding to the trimeric spike. The heterodimeric forms that conjugate two Nbs of unique, non-overlapping epitopes, through a flexible linker of 12 residues.

**Figure 5.**
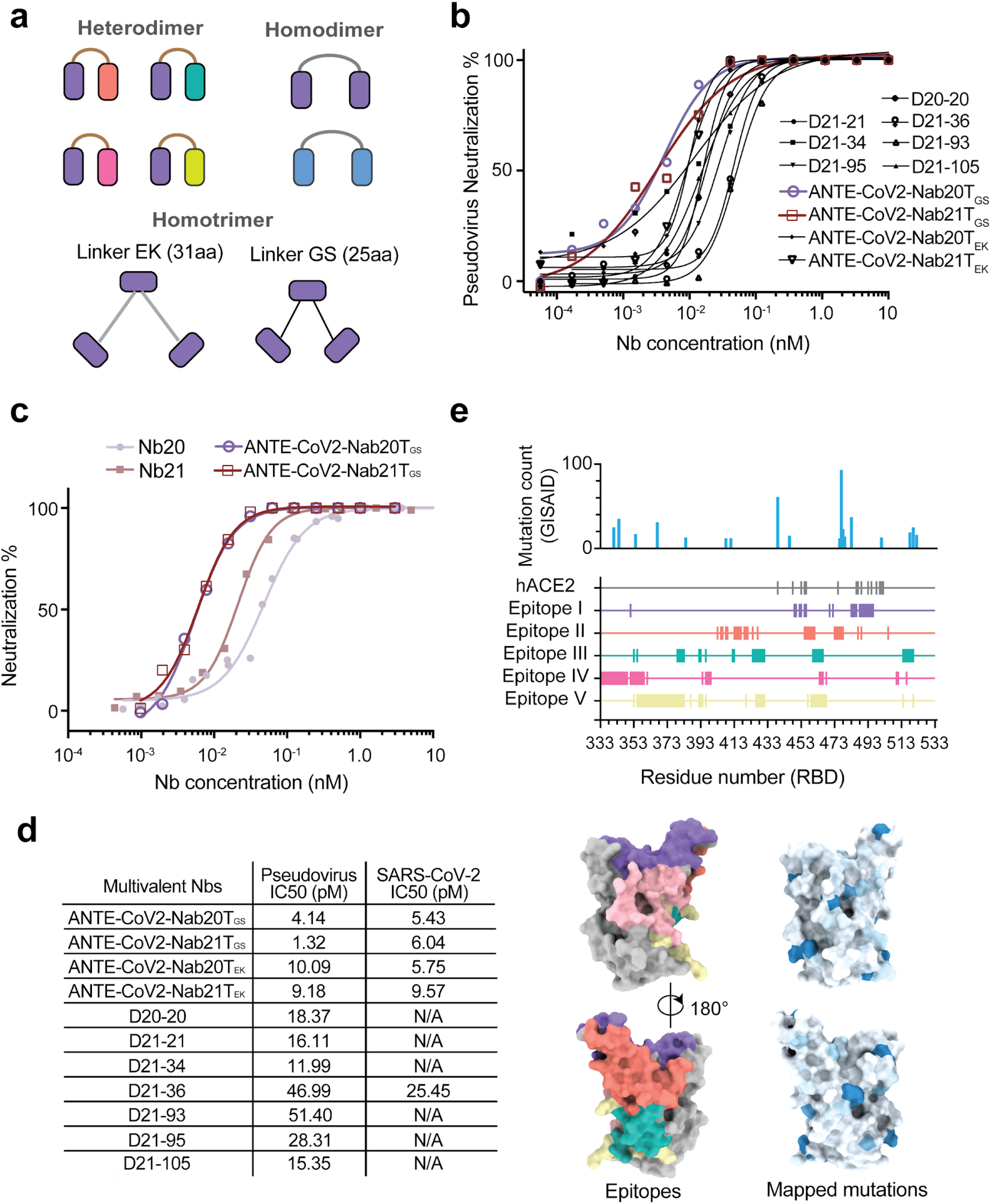
Development of multivalent Nb cocktails for highly efficient SARS-CoV-2 neutralization. **Fig 5a**: Schematics of the cocktail design **Fig 5b**: Pseudotyped SARS-CoV-2 neutralization assay of multivalent Nbs. The average neutralization percentage of each data point was shown (n=2). **Fig 5c**: SARS-CoV-2 PRNT of monomeric and trimeric forms of Nbs 20 and 21. The average neutralization percentage of each data point was shown (n=2 for the trimers, n=4 for the monomers). **Fig 5d**: A summary table of the neutralization potency measurements of the multivalent Nbs. N/A: not tested. **Fig 5e**: Mapping mutations to localization of Nb epitopes on the RBD. The x-axis corresponds to the RBD residue numbers (333 to 533). Rows in different colors represent different epitope residues. Epitope I: 351, 449-450, 452-453, 455-456, 470, 472, 483-486, 488-496; Epitope II: 403, 405-406, 408,409, 413-417, 419-421, 424, 427, 455-461, 473-478, 487, 489, 505; Epitope III: 53, 355, 379-383, 392-393, 396, 412-413, 424-431, 460-466, 514-520; Epitope IV: 333-349, 351-359, 361, 394, 396-399, 464-466, 468, 510-511, 516; Epitope V: 353, 355-383, 387, 392-394, 396, 420, 426-431, 457,459-468, 514, 520

We synthesized a variety of constructs and tested their neutralization potency. We found up to ~30 fold improvement for the homotrimeric constructs of Nb21_3_ (IC50 =1.3 pM) and Nb20_3_ (IC50 =3 pM) compared to the respective monomeric form by the pseudovirus luciferase assay (**Fig 5b, 5d**). Similar results were obtained from the SARS-CoV-2 PRNT (**Fig 5c, 5d, S10a)**. The improvements are likely greater than these values indicate, as the measured values may reflect the assay’s lower detection limits. For the heterodimeric constructs, up to a 4-fold increase of potency (i.e., Nb21-Nb34) was observed. Importantly, the multivalent constructs retained outstanding physicochemical properties of the monomeric Nbs, including high solubility, yield, and thermostability (**Fig S9**). They remained fully active after standard lyophilization and nebulization (**Methods**, **Fig S10b-e**), indicating the outstanding stability and flexibility of administration. The majority of the RBD mutations observed in GISAID {Shu, 2017 #68} are very low in frequency (<0.0025). Therefore, the probability of mutational escape with a cocktail consisting of 2-3 Nbs covering different epitopes is extremely low (**Fig 5e**) {Hansen, 2020 #8}.

## Discussion

The development of effective, safe, and inexpensive vaccines and therapeutics are critical to end the COVID-19 pandemic. Here, *in vivo* (camelid) antibody affinity maturation followed by advanced proteomics {Xiang, 2020 #67} enabled rapid identification of a large repertoire of diverse, high-affinity RBD Nbs for the neutralization of SARS-CoV-2. The majority of the high-affinity Nbs efficiently neutralize SARS-CoV-2 and some elite Nbs in their monomeric forms can inhibit the viral infection at single-digit to sub-ng/ml concentrations.

We identified multiple neutralization epitopes through the integration of biophysics, structural proteomics, modeling, and X-ray crystallography. We have gained novel insight into the mechanisms by which Nbs target the RBD with femtomolar affinity to achieve remarkable neutralization potency of a low-passage, clinical isolate of SARS-CoV-2. Structural analysis revealed that the hACE2 binding site correlates with immunogenicity and neutralization. While the most potent Nbs inhibit the virus by high-affinity binding to the hACE2 binding site, we also observed other neutralization mechanisms through non-hACE2 epitopes.

A preprint reported an extensively bioengineered, homotrimeric Nb construct reaching antiviral activity comparable to our single monomeric Nb20 {Schoof, 2020 #32}. Here, we have developed a collection of novel multivalent Nb constructs with outstanding stability and neutralization potency at double-digit pg/ml. To our knowledge, this represents the most potent biotherapeutics for SARS-CoV-2 available to date. The use of multivalent, multi-epitope Nb cocktails may prevent virus escape {Baum, 2020 #26;Bar-On, 2018 #44;Marovich, 2020 #45}. Flexible and efficient administration, such as direct inhalation could be used to improve antiviral drug efficacy and minimize the dose, cost, and potential toxicity for clinical applications. The high sequence similarity between Nbs and IgGs may restrain the immunogenicity {Jovcevska, 2020 #58}. For intravenous drug delivery, it is possible to fuse our antiviral Nbs with the albumin-Nb constructs {Shen, 2020 #31} t already developed to improve the *in vivo* half-lives. These Nbs can also be applied as rapid point-of-care diagnostics due to the high stability, specificity, and low cost of manufacturing. We envision that these high-quality Nb agents will contribute to curbing the current pandemic.

## Acknowledgments

We thank the staff at the GM/CA of APS in the Argonne National Laboratory (US) for their assistance with X-ray diffraction data collection. We thank Prof. Zhiyi Wei at the Southern University of Science and Technology for help with crystal structure determination. This work was supported by the University of Pittsburgh School of Medicine (Y.S.), CTSI pilot fund (Y.S.), The University of Pittsburgh and the Center for Vaccine Research (WPD), NIH grant R35GM128641 (C.Z.), ISF 1466/18 (D.S.), and Israeli Ministry of Science and Technology (D.S.).

## Contributions

Y.S. and D.S conceived the study. Y.X. performed most of the experiments. S.N. performed the PRNT SARS-CoV-2 neutralization assay. Z.X. produced the multivalent Nbs and performed thermostability measurements. C.Z. determined the X-ray structure. Y.X., Y.S., D.S., C.Z., Z.S., S.N., and P.D. analyzed the data. Y.S. cheerled the study and drafted the manuscript. All authors edited the manuscript.

## Supplemental Materials

**Figure S1.**
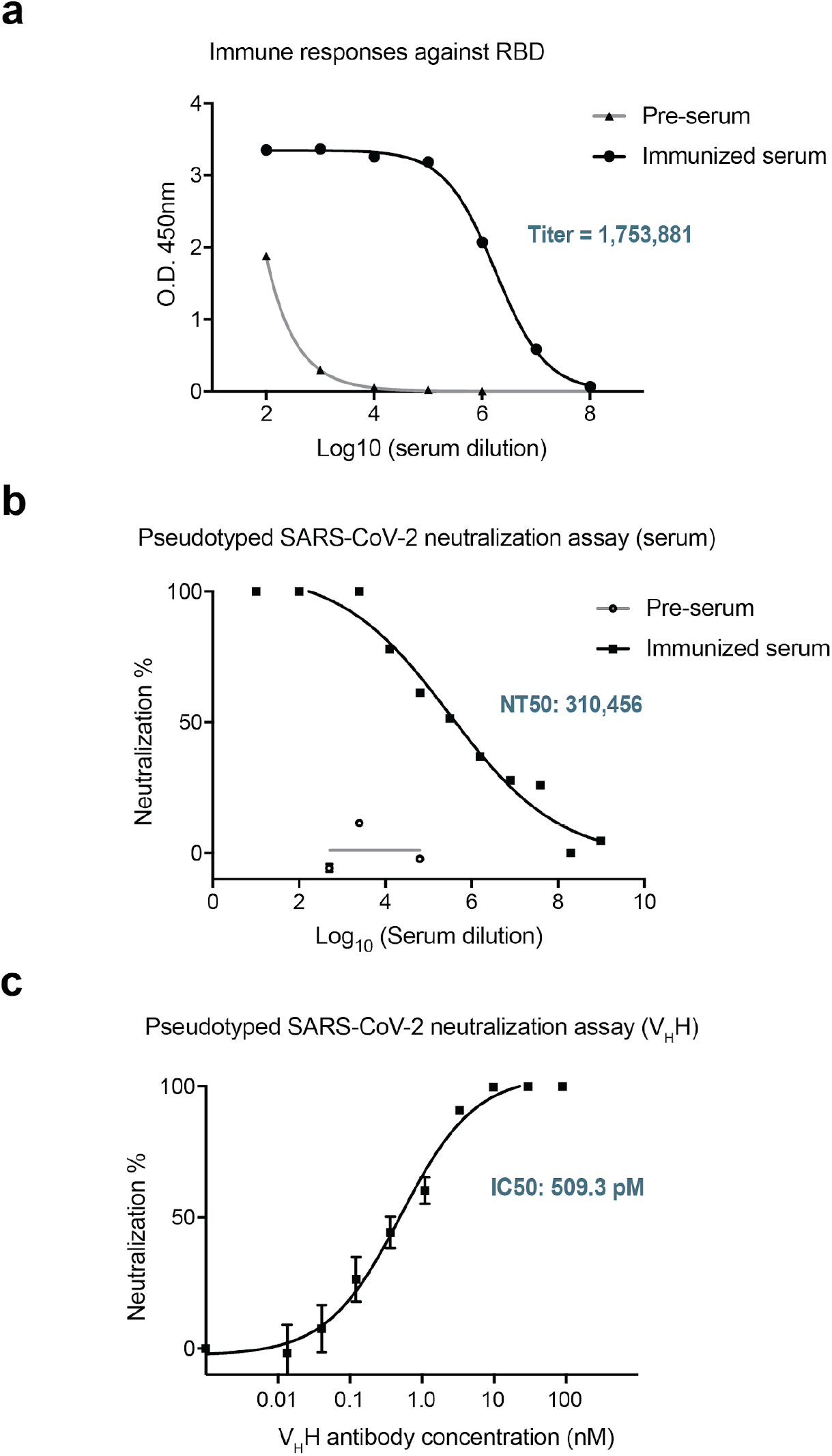
Development of RBD-specific Nbs for potent SARS-CoV-2 neutralization. S1a: Detection of strong and specific serologic activities after immunization of SARS-CoV-2 _RBD._ S1b: Neutralization potency of the immunized camelid’s serum against pseudotyped SARS-CoV-2 (luciferase) S1c: Neutralization potency of Nbs against pseudotyped SARS-CoV-2 (luciferase).

**Figure S2.**
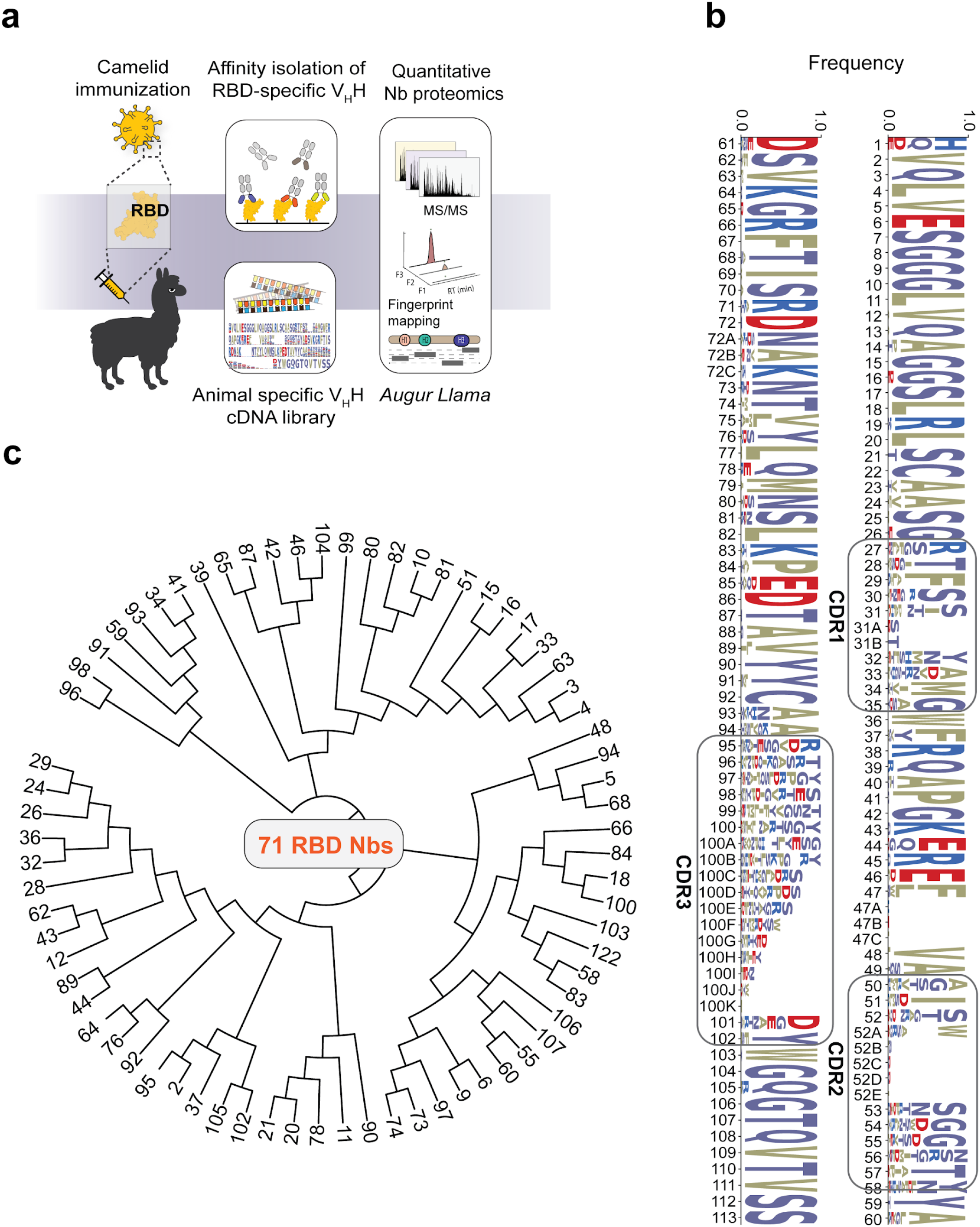
Identification of high-affinity RBD-Nbs. S2a: High-affinity RBD-specific Nb identification by camelid immunization and quantitative Nb proteomics. Briefly, a camelid was immunized with the RBD protein. After immunization, RBD-specific V_H_H antibodies were isolated from the serum, and analyzed by quantitative proteomics to identify the high-affinity, RBD-specific Nbs (see Methods). A V_H_H (Nb) cDNA library from the plasma B cells of the immunized camelid was generated to facilitate proteomic analysis. S2b: Sequence logo of 71 RBD Nbs. Each Nb has a unique CDR3 sequence. The amino acid occurrence at each position is shown. CDR: complementarity determining region. FR: framework. S2c: The phylogenetic tree of the Nbs constructed by the maximum likelihood model.

**Figure S3.**
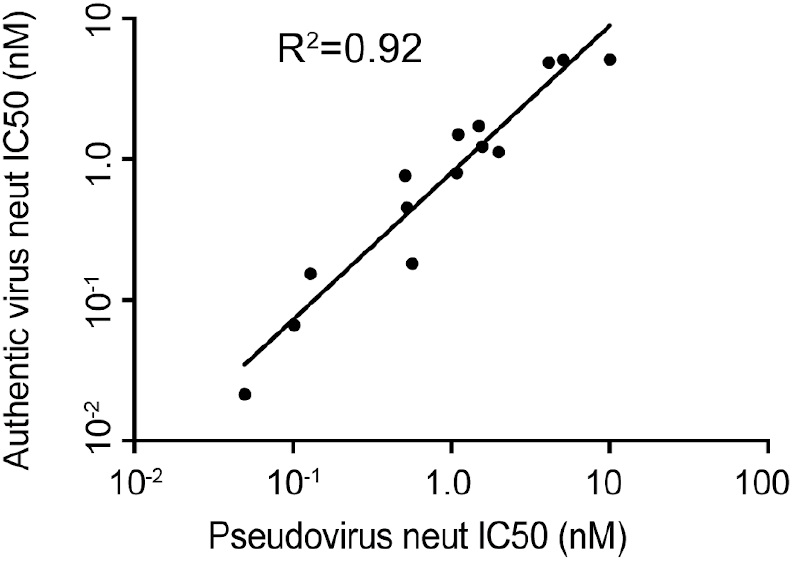
Correlation analysis of 18 highly potent SARS-CoV-2 neutralizing Nbs. A plot showing a linear correlation of Nb neutralization IC50s between the pseudotyped virus neutralization assay and the SARS-CoV-2 PRNT.

**Figure S4.**
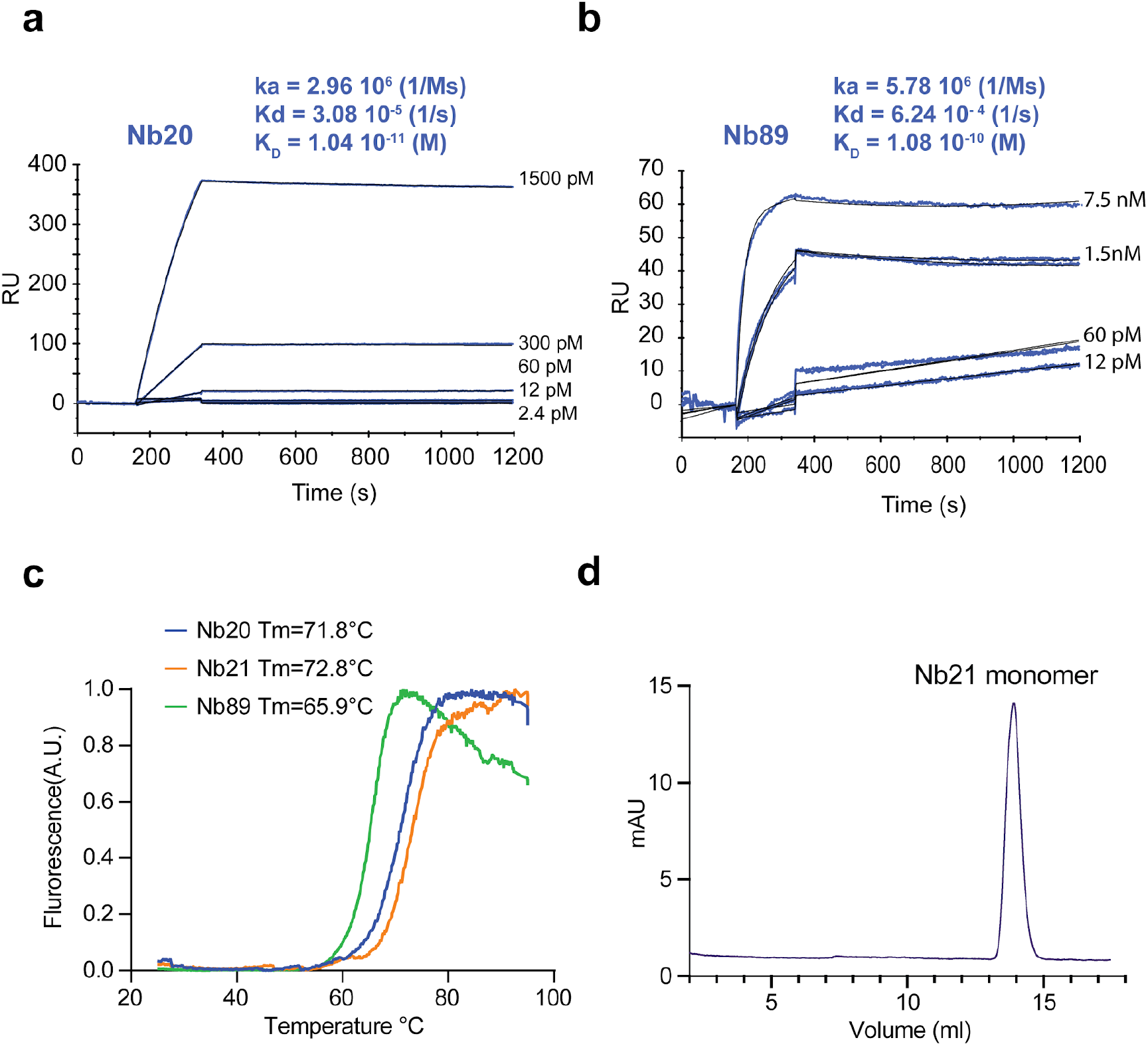
Biophysical analysis of the outstanding neutralizing Nbs. S4a-b: Binding kinetics of Nbs 20 and 89 by SPR. S4c: Thermostability analysis of Nbs 20, 21, and 89. The values represent the average thermostability (Tm, °C) based on three replicates. The standard deviations (SD) of the measurements are 0.17, 0.93, and 0.8 °C for Nbs 20, 21, and 89. S4d: Stability analysis of Nb21 by SEC. Purified recombinant Nb21 was stored at room temperature for ~ 6 weeks before subject to SEC analysis. The dominant peak represents Nb 21 monomer.

**Figure S5.**
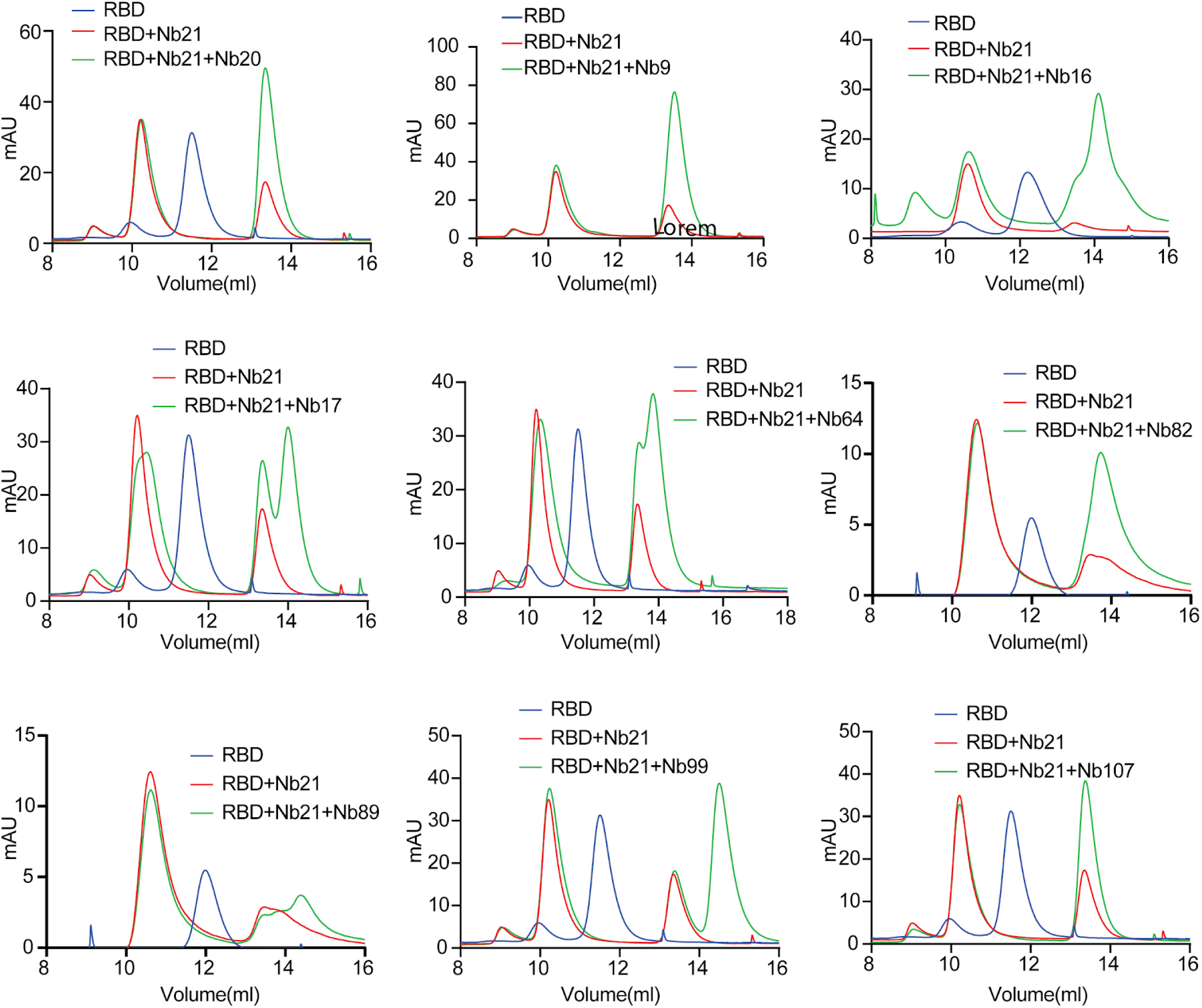
SEC analysis of RBD-Nb complexes. The SEC profiles of RBD-Nb complexes showing 9 Nbs that have overlapping epitopes with Nb 21.

**Figure S6.**
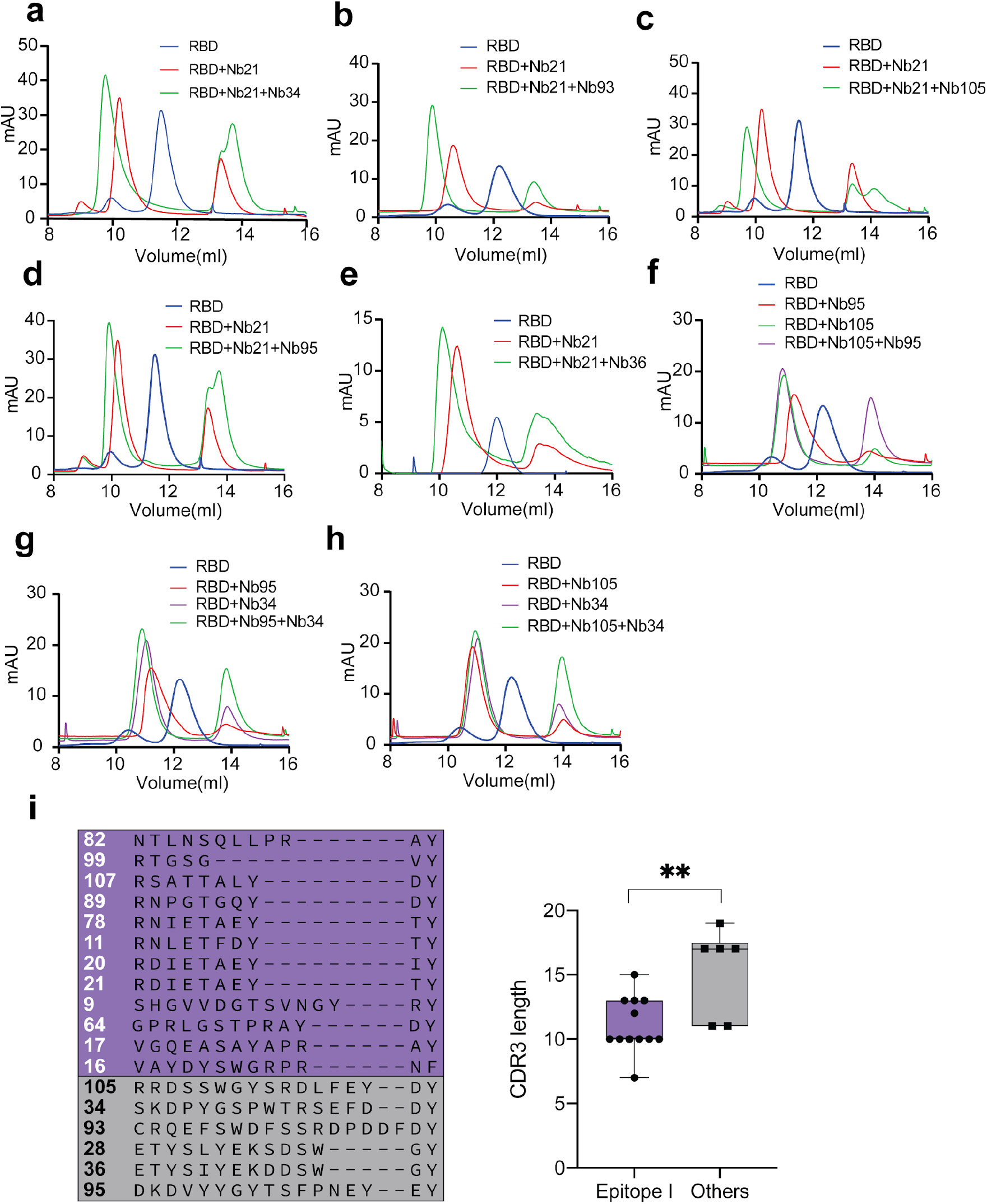
SEC analysis of RBD-Nb complexes. S6a-h: The SEC profiles of RBD-Nb complexes showing five Nbs that have unique and non-overlapping epitopes with Nb21. S6i: Sequence alignment of the CDR3s of 18 highly potent neutralizing Nbs and CDR3 lengths comparing Nbs from epitope I and others.

**Figure S7.**
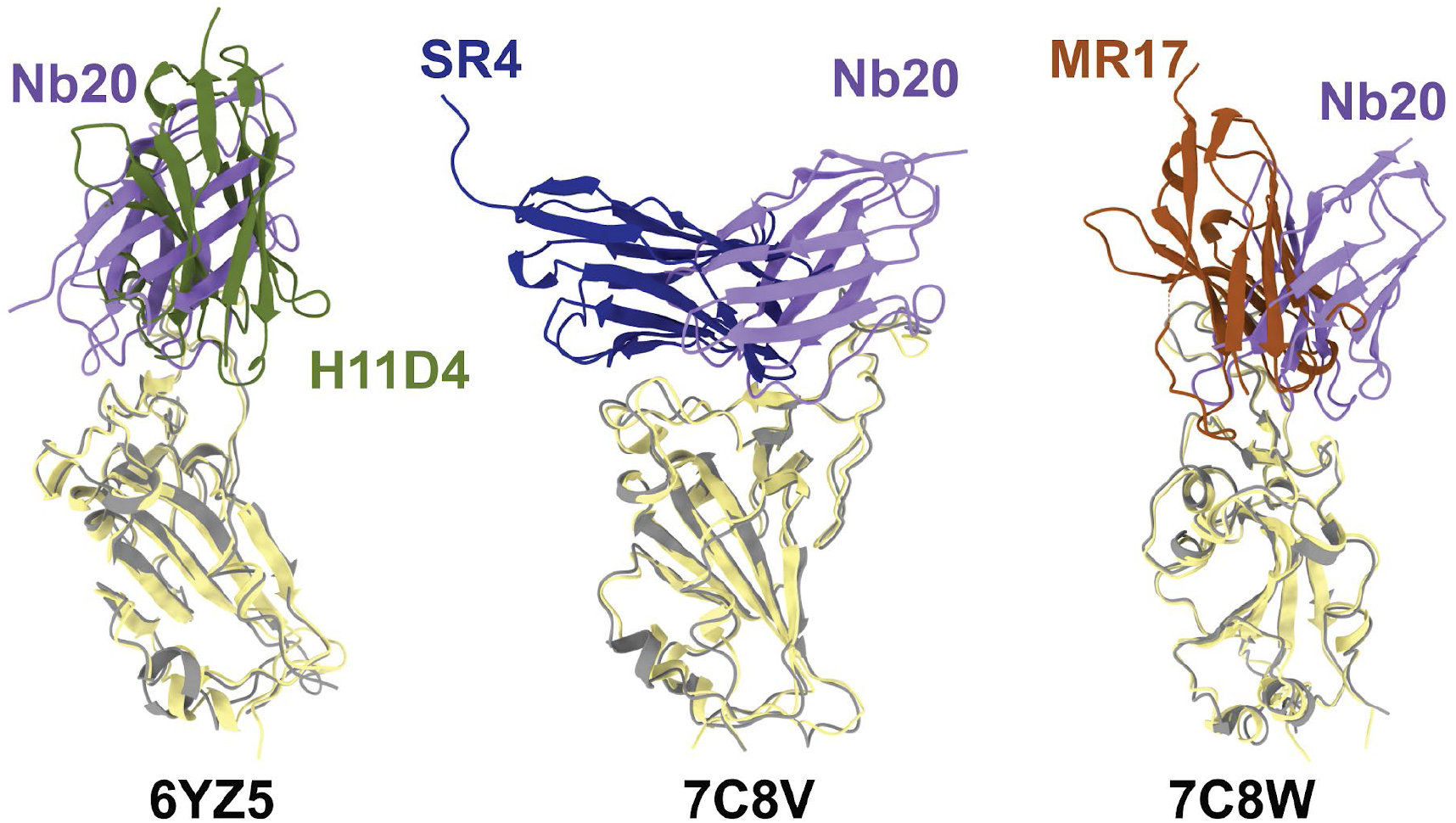
Structural comparisons of Nb20 with published RBD Nb structures. Overlays of Nb20 (purple ribbon) and three other RBD-Nbs (PDBs 6YZ5, 7C8V, and 7C8W) in complex with RBD (yellow/grey ribbon).

**Figure S8.**
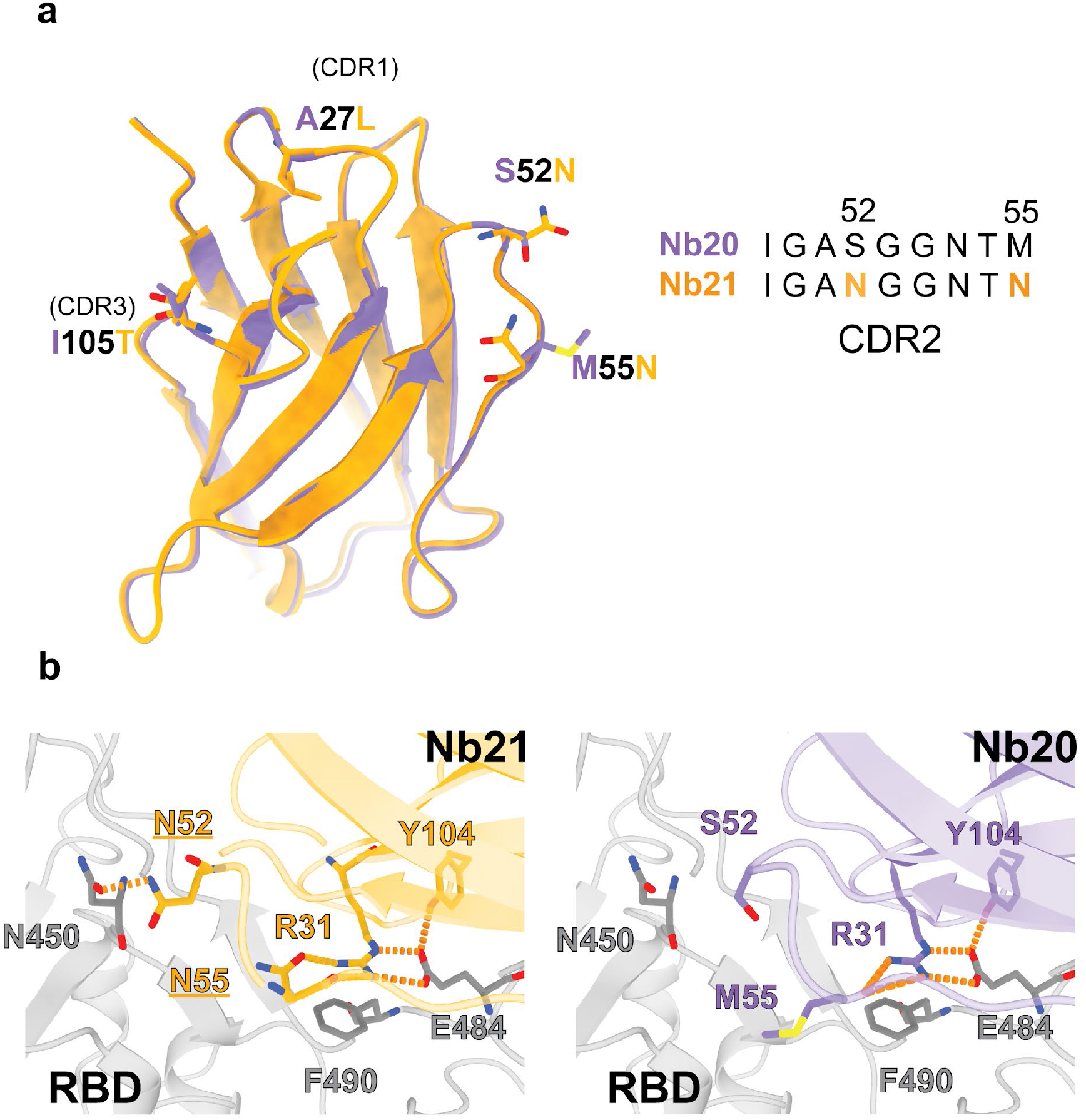
Modeling of Nb 21-RBD interaction based on the crystal structure of Nb 20-RBD complex. S8a: Alignment of Nb21 with Nb20. The four residue differences were shown. S8b: Zoom-in views showing the addition of new polar interaction between N52 (Nb21) and N450 (RBD). The model of Nb21 is superimposed based on the crystal structure of Nb20.

**Figure S9.**
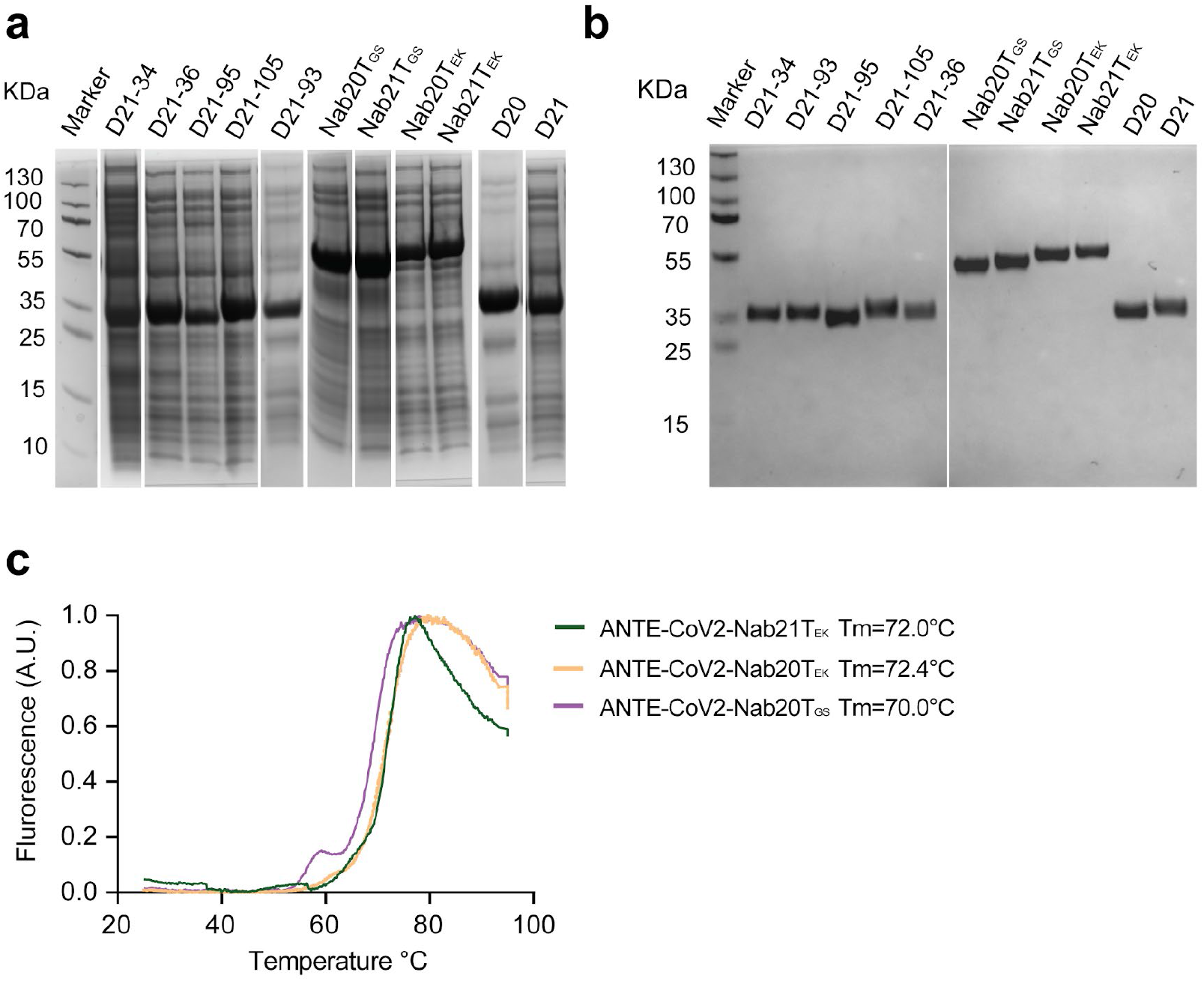
The biophysical properties of multivalent Nbs. S9a: The expression levels of multivalent Nbs from *E.coli* whole cell lysate. S9b: A SDS-PAGE gel picture showing the purification of the multivalent Nbs. S9b: Thermostability analysis of ANTE-CoV2-Nab21T_EK_, ANTE-CoV2-Nab20T_EK_, and ANTE-CoV2-Nab20T_GS_. The values represent the average thermostability (Tm, °C) based on three replicates. The standard deviations of the measurement are 0.6, 0.27, and 0.72°C, respectively.

**Figure S10.**
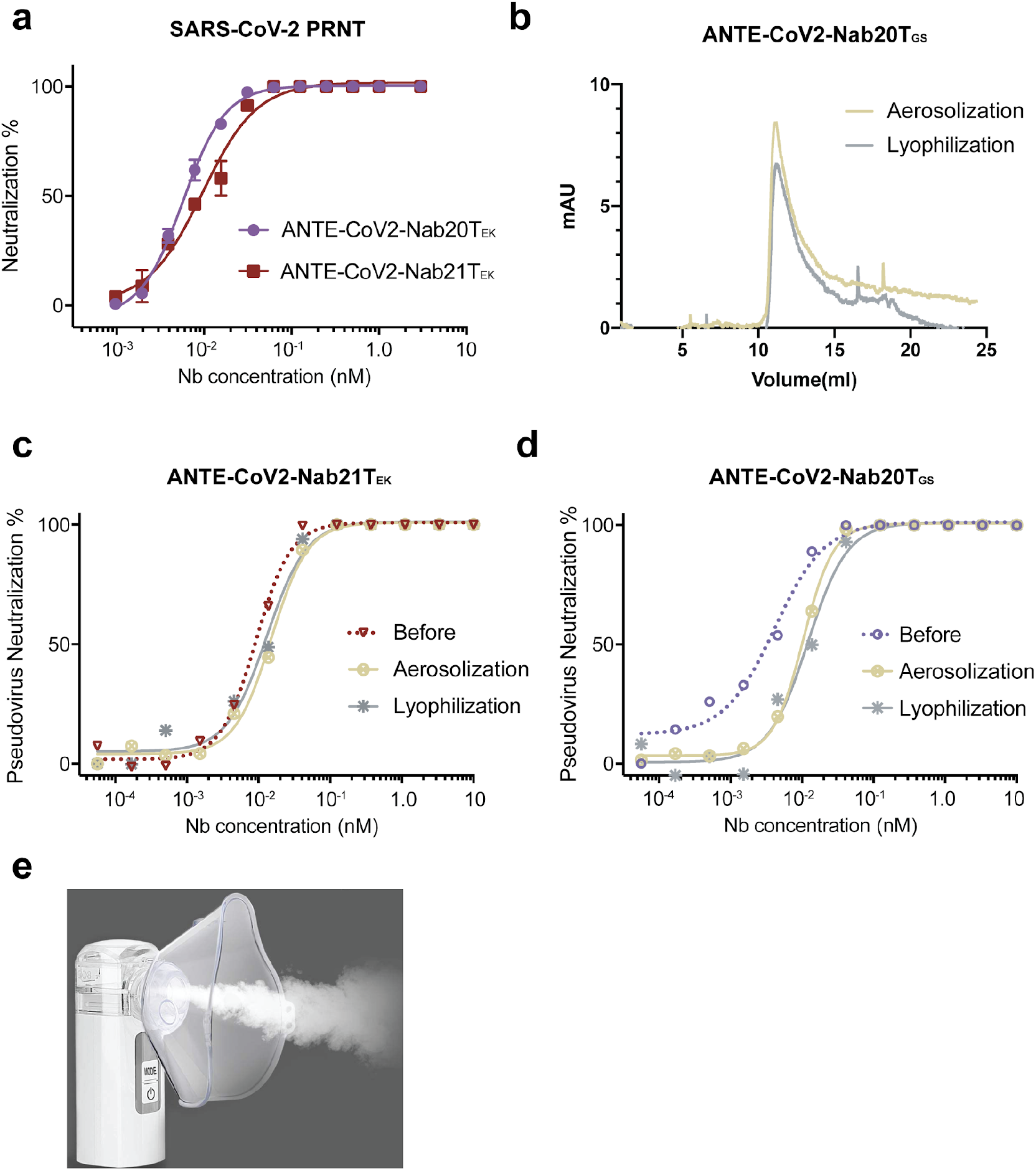
Stability test of the multivalent Nbs. S10a: SARS-CoV-2 PRNT of trimeric forms of Nbs 20 and 21 with the EK linker. The average neutralization % and the standard deviation of each data point were shown (n=2). S10b: The example SEC analyses of ANTE-CoV2-Nab20T_GS_ after lyophilization and aerosolization. S10c: Pseudotyped SARS-CoV-2 neutralization assay using ANTE-CoV2-Nab21T_EK_ before and after lyophilization and aerosolization. S10d: Pseudotyped SARS-CoV-2 neutralization assay using ANTE-CoV2-Nab20T_GS_ before and after lyophilization and aerosolization. S10e: Portable mesh nebulizer (producing ≤ 5μm aerosol particles) used in the study.

**Supplemental Database.** Amino acid sequences of the 71 RBD-Nbs and 11 multivalent Nbs.

**Supplemental Table 1.** Summary of the biophysical, biochemical and functional properties of the monomeric RBD-Nbs and the multivalent constructs.

**Supplemental Table 2.** Summary of the cross-links of the Nbs from different epitopes.

**Supplemental Table 3.** X-ray diffraction data collection and refinement statistics.

|  | SARS-CoV-2 RBD with Nb20<br>(PDB code: 7JVB) |
| --- | --- |
| **Data collection** |  |
| Space group | $P4_12_12$ |
| Cell dimensions |  |
| *a*, *b*, *c* (Å) | 70.716, 70.716, 435.037 |
| $\alpha$, $\beta$, $\gamma$ (°) | 90, 90, 90 |
| Resolution (Å) | 40-3.3 (3.36-3.30) |
| $R_{\text{merge}}$ (%)<sup>a</sup> | 15.6 (59.1) |
| $I/\sigma(I)$ | 7.8 (1.1) |
| $CC_{1/2}$<sup>b</sup> | 1(0.811) |
| Completeness (%) | 97.1 (81.4) |
| Redundancy | 6.2 (3.4) |
| **Refinement** |  |
| Resolution (Å) | 38.96-3.287 (3.405-3.287) |
| No. reflections | 17336 (1377) |
| $R_{\text{work}} / R_{\text{free}}$ (%)<sup>c</sup> | 28.7 (38.2) / 32.2 (43.0) |
| No. atoms |  |
| Protein | 4728 |
| Ligand/ion | 20 |
| Water | 0 |
| *B* factors |  |
| Protein | 133.33 |
| Ligand/ion | 120.40 |
| Water | N/A |
| R.m.s. deviations |  |
| Bond lengths (Å) | 0.005 |
| Bond angles (°) | 1.57 |
| Ramachandran analysis |  |
| Favored region (%) | 92.16 |
| Allowed region (%) | 7.84 |
| Outliers (%) | 0 |

The numbers in parentheses represent values for the highest resolution shell.
<sup>a</sup> $R_{\text{merge}} = \sum |I_i - I_m| / \sum I_i$ , where $I_i$ is the intensity of the measured reflection and $I_m$ is the mean intensity of all symmetry-related reflections.
<sup>b</sup> $CC_{1/2}$ is the correlation coefficient of the half datasets.
<sup>c</sup> $R_{\text{work}} = \sum ||F_{\text{obs}}| - |F_{\text{calc}}|| / \sum |F_{\text{obs}}|$ , where $F_{\text{obs}}$ and $F_{\text{calc}}$ are observed and calculated structure factors.
$R_{\text{free}} = \sum T ||F_{\text{obs}}| - |F_{\text{calc}}|| / \sum T |F_{\text{obs}}|$ , where $T$ is a test data set of about 5% of the total reflections randomly chosen and set aside prior to refinement.

## Materials and methods

### Camelid immunization and proteomic identification of high-affinity RBD-Nbs

A male Llama “Wally” was immunized with an RBD-Fc fusion protein (Acro Biosystems, Cat#SPD-c5255). The bleeds from the animal were collected after a 55-day immunization protocol (Capralogics, Inc.). All the above procedures were performed by following the IACUC protocol. ~ 1 ×10^9^ peripheral mononuclear cells were isolated from 350 ml immunized blood using Ficoll gradient (Sigma). The mRNA was purified from the mononuclear cells using an RNeasy kit (Qiagen) and was reverse-transcribed into cDNA by the Maxima™ H Minus cDNA Synthesis kit (Thermo). The VHH genes were PCR amplified, and the P5 and P7 adapters were added with the index before sequencing. Next-generation sequencing (NGS) of the VHH repertoire was performed by Illumina MiSeq with the 300 bp paired-end model.

For proteomic analysis of RBD-specific Nbs, plasma was first purified from the immunized blood by the Ficoll gradient (Sigma). V_H_H antibodies were then isolated from the plasma by a two-step purification protocol using protein G and protein A sepharose beads (Marvelgent) {Fridy, 2014 #46}. RBD-specific V_H_H antibodies were affinity isolated and subsequentially eluted by either increasing stringency of high pH buffer or salt. All the eluted V_H_Hs were neutralized and dialyzed into 1x DPBS before the quantitative proteomics analysis. RBD-specific V_H_H antibodies were reduced, alkylated and in-solution digested using either trypsin or chymotrypsin {Xiang, 2020 #67}. After proteolysis, the peptide mixtures were desalted by self-packed stage-tips or Sep-Pak C18 columns (Waters) and analyzed with a nano-LC 1200 that is coupled online with a Q Exactive™ HF-X Hybrid Quadrupole Orbitrap™ mass spectrometer (Thermo Fisher). Proteomic analysis was performed as previously described and by using the Augur Llama-a dedicated software that we developed to facilitate reliable identification, label-free quantification, and classification of high-affinity Nbs {Xiang, 2020 #67}. This analysis led to thousands of RBD-specific, high-affinity Nb candidates that belong to ~ 350 unique CDR3 families. From these, we selected 109 Nb sequences with unique CDR3s for DNA synthesis and characterizations.

### Nb DNA synthesis and cloning

The monomeric Nb genes and the homotrimeric Nbs20 and 21 with the (GGGGS)_5_ linkers were codon-optimized and synthesized (Synbio). Nb DNA sequences were cloned into a pET-21b(+) vector or pET-22b(+) vector using EcoRI and HindIII or BamHI and XhoI restriction sites. To produce a heterodimeric Nb such as Nb21-Nb34, the DNA fragment of the monomeric Nb34 was first PCR amplified from the pET-21b(+) vector to introduce a linker sequence and two restriction sites of XhoI and HindIII that facilitate cloning. The PCR fragment was then inserted into the Nb21 pET-21b(+)vector to produce the heterodimer Nb21-(GGGGS)_2_-Nb34. Homodimers or homotrimers with the EK linker were produced in-house. For the in-house generated constructs, a flexible linker sequence of EGKSSGSGSESKSTGGGGSEGKSSGSGSESKST introduced using the following two oligos:

CCGCTCGAGTGCTGCGGCCGCGGTGCTTTTGCTTTCGCCGCTACCGCTGCTTTTACCTTCGCTGCCACC, and
CCCAAGCTTGAAGGTAAAAGCAGCGGTAGCGGCGAAAGCAAAAGCACCGGTGGCGGTGGCAGCGAAGGT (Integrated DNA Technologies).

Briefly, after digestion using XhoI/HindIII restriction sites, the linker was inserted into pET21b(+)_Nb21 or pET21b(+)_Nb20 vector. For simplicity, here we used the Nb21 trimer for illustration of the cloning strategy. To shuffle the second copy of an Nb, we amplified Nb 21 from the pET21b(+) vector and introduced XhoI/NotI restriction sites. After digestion, the XhoI/NotI Nb fragment was inserted into the Nb_linker vector to produce an Nb homodimer. The following DNA oligos were used to facilitate the cloning of the second copy of the linker in which EcoRI and BamHI sites were introduced:

CCGGAATTCGAGGTGCTTTTGCTTTCGCCGCTACCGCTGCTTTTACCTTCGCTGCCACCGCCACCGGTGC, and
CGCGGATCCGGGTTCGAGCTCGGAAGGTAAAAGCAGCGGTAGCGGCGAAAGCAAAAGCACCGGTGGCGG (Integrated DNA Technologies).

After digestion, the second linker was then inserted into the homodimeric pET21b(+)_Nb21 vector. The third copy of the Nb21 gene was PCR amplified from the vector and inserted into the homodimeric pET21b(+)_Nb21 construct using BamHI/SacI sites. All the DNA constructs were verified by sanger sequencing.

### Expression and purification of proteins

Nb DNA constructs were transformed into BL21(DE3) cells and plated on Agar with 50 μg/ml ampicillin at 37 °C overnight. Cells were cultured in an LB broth to reach an O.D. of ~ 0.5-0.6 before IPTG (0.5 mM) induction at 16°C overnight. Cells were then harvested, sonicated, and lysed on ice with a lysis buffer (1xPBS, 150 mM NaCl, 0.2% TX-100 with protease inhibitor). After cell lysis, protein extracts were collected by centrifugation at 15,000 X g for 10 mins and the his-tagged Nbs were purified by Cobalt resin and natively eluted by imidazole buffer (Thermo). Eluted Nbs were subsequently dialyzed in a dialysis buffer (e.g., 1x DPBS, pH 7.4). For periplasmic preparation of Nbs, cell pellets were resuspended in the TES buffer (0.1M Tris-HCl, pH 8.0; 0.25mM EDTA, pH 8.0; 0.25M Sucrose) and incubated on ice for 30 min. The supernatants were collected by centrifugation and subsequently dialyzed to DPBS. The resulting Nbs were then purified by Cobalt resin as described above.

The RBD (residues 319-541) of the SARS-Cov-2 S protein was expressed as a secreted protein in *Spodoptera frugiperda* Sf9 cells (Expression Systems) using the Bac-to-bac baculovirus method (Invitrogen). To facilitate protein purification, a FLAG-tag and an 8 × His-tag were fused to its N terminus, and a tobacco etch virus (TEV) protease cleavage site was introduced between the His-tag and RBD. Cells were infected with baculovirus and incubated at 27 °C for 60 h before harvesting. The conditioned media was added with 20 mM Tris pH 7.5 and incubated at RT for 1 h in the presence of 1 mM NiSO_4_ and 5mM CaCl_2_. The supernatant was collected by centrifugation at 25,000 g for 30 min and then incubated with Nickel-NTA agarose resin (Clontech) overnight at 4 °C. After washing with buffer containing 20 mM Hepes pH 7.5, 200 mM NaCl, and 50mM imidazole, the RBD protein was eluted with the same buffer containing 400 mM Imidazole. Eluted protein was treated by TEV protease overnight to remove extra tags and further purified by size exclusion chromatography using the Superdex 75 column (Fisher) with a buffer containing 20 mM Hepes pH 7.5 and 150 mM NaCl. To obtain RBD and Nb20 complex, purified RBD was mixed with purified Nb20 in a molar ratio of 1: 1.5 and then incubated on ice for 2 hours. The complex was further purified using the Superdex 75 column with a buffer containing 20 mM Hepes pH 7.5 and 150 mM NaCl. Purified RBD-Nb20 complex was concentrated to 10-15 mg/ml for crystallization.

### ELISA (Enzyme-linked immunosorbent assay)

Indirect ELISA was carried out to measure the relative affinities of Nbs. RBD was coated onto a 96-well ELISA plate (R&D system) at two ng/well in coating buffer (15 mM sodium carbonate, 35 mM sodium bicarbonate, pH 9.6) overnight at 4°C and was blocked with a blocking buffer (DPBS, 0.05% Tween 20, 5% milk) at room temperature for 2 hrs. Nbs were serially 10X diluted in the blocking buffer, starting from 1 μM to 0.1 pM, and 100 μl of each concentration was incubated with RBD-coated plates for 2 hrs. HRP-conjugated secondary antibodies against T7-tag (Thermo) were diluted 1:7500 and incubated with the well for 1 hr at room temperature. After PBST (DPBS, 0.05% Tween 20) washes, the samples were further incubated under dark with freshly prepared w3,3′,5,5′-Tetramethylbenzidine (TMB) substrate for 10 mins to develop the signals. After the STOP solution (R&D system), the plates were read at multiple wavelengths (the optical density at 550 nm wavelength subtracted from the density at 450 nm) on a plate reader (Multiskan GO, Thermo Fisher). A non-binder was defined if any of the following two criteria were met: i) The ELISA signal was under detected at one μM concentration. ii) The ELISA signal could only be detected at a concentration of 1 μM and was under detected at 0.1 μM concentration. The raw data was processed by Prism 7 (GraphPad) to fit into a 4PL curve and to calculate logIC50.

### Pseudotyped SARS-CoV-2 neutralization assay

The 293T-hsACE2 stable cell line (Cat# C-HA101) and the pseudotyped SARS-CoV-2 (Wuhan-Hu-1 strain) particles with GFP (Cat# RVP-701G, Lot#CG-113A) or firefly luciferase (Cat# RVP-701L, Lot# CL109A, and CL-114A) reporters were purchased from the Integral Molecular. The neutralization assay was carried out according to the manufacturers’ protocols. In brief, 10-fold serially diluted Nbs were incubated with the pseudotyped SARS-CoV-2-GFP for 1 hr at 37 °C for screening, while 3- or 5-fold serially diluted Nbs / immunized serum / immunized V_H_H mixture was incubated with the pseudotyped SARS-CoV-2-luciferase for accurate measurements. At least eight concentrations were tested for each Nb. Pseudovirus in culture media without Nbs was used as a negative control. The mixtures were then incubated with 293T-hsACE2 cells at 2.5×10e5 cells/ml in the 96-well plates. The infection took ~72 hrs at 37 °C with 5% CO_2_. The GFP signals (ex488/em530) were read using the Tecan Spark 20M with auto-optimal settings, while the luciferase signal was measured using the *Renilla*-Glo luciferase assay system (Promega, Cat# E2720) with the luminometer at 1 ms integration time. The obtained relative fluorescent/luminescence signals (RFU/RLU) from the negative control wells were normalized and used to calculate neutralization percentage for each concentration. For SARS-CoV-2-GFP screening, the 49 tested Nbs were divided into 6 groups based on their lowest tested concentration of 100% neutralization. For SARS-CoV-2-luciferase, data was processed by Prism7 (GraphPad) to fit into a 4PL curve and to calculate the logIC50 (half-maximal inhibitory concentration).

### SARS-CoV-2 Munich plaque reduction neutralization test (PRNT)

Nbs were diluted in a 2- or 3-fold series in Opti-MEM (Thermofisher). Each Nb dilution (110 μl) was mixed with 110 μl of SARS-CoV-2 (Munich strain) containing 100 plaque-forming units (p.f.u.) of the virus in Opti-MEM. The serum–virus mixes (220 μl total) were incubated at 37 °C for 1 h, after which they were added dropwise onto confluent Vero E6 cell monolayers in the six-well plates. After incubation at 37 °C, 5% (v/v) CO_2_ for 1 h, 2 ml of 0.1% (w/v) immunodiffusion agarose (MP Biomedicals) in Dulbecco's modified eagle medium (DMEM) (Thermofisher) with 10% (v/v) FBS and 1× pen-strep was added to each well. The cells were incubated at 37 °C, 5% CO_2_ for 72 h. The agarose overlay was removed and the cell monolayer was fixed with 1 ml/well formaldehyde (Fisher) for 20 min at room temperature. Fixative was discarded and 1 ml/well of 1% (w/v) crystal violet in 10% (v/v) methanol was added. Plates were incubated at room temperature for 20 min and rinsed thoroughly with water. Plaques were then enumerated and the 50% plaque reduction neutralization titer (PRNT50) was calculated. A validated SARS-CoV-2 antibody-negative human serum control, a validated NIBSC SARS-CoV-2 plasma control, was obtained from the National Institute for Biological Standards and Control, UK) and an uninfected cells control were also performed to ensure that virus neutralization by antibodies was specific.

### Surface plasmon resonance (SPR)

Surface plasmon resonance (Biacore 3000, GE) was used to measure Nb affinities. Briefly, recombinant Nb was immobilized to the flow channels of an activated CM5 sensor-chip. Nb was diluted to 10 μg/ml in 10 mM sodium acetate, pH 5.5, and injected into the SPR system at 5 μl/min for 420 s. The surface was then blocked by 1 M ethanolamine-HCl (pH 8.5). For RBD analyte, a series of dilution (spanning ~ 1,000-fold concentration range) was injected in duplicate, with HBS-EP running buffer (GE-Healthcare) at a flow rate of 30 μl/min for 180 s, followed by a dissociation time of 20 mins. Between each injection, the sensor chip surface was regenerated twice with 6M Guanidine-HCl at a flow rate of 40-50 μl/min for 30 s - 1 min. Binding sensorgrams for each Nb were processed and analyzed using BIA evaluation by fitting with the 1:1 Langmuir model.

### Phylogenetic tree analysis and sequence logo

Sequences were first aligned and numbered according to Martin’s numbering scheme by ANARCI {Dunbar, 2016 #47}. The phylogenetic tree was constructed from aligned sequences by Molecular Evolutionary Genetics Analysis(MEGA){Kumar, 2018 #48} using the Maximum Likelihood method. The sequence logo was plotted from aligned sequences by logomaker{Tareen, 2020 #49}.

### Epitope screening by size exclusion chromatography (SEC)

Recombinant RBD and Nb proteins were mixed at a ratio of 1:1 (w:w) and incubated at 4°C for 1 hr. The complexes were analyzed by the SEC (Superdex75, GE LifeSciences) at a low rate of 0.4 ml/min for 1 hr using a running buffer of 20 mM HEPES, 150 mM NaCl, pH 7.4. Protein signals were detected by ultraviolet light absorbance at 280 nm.

### Chemical cross-linking and mass spectrometry (CXMS)

Recombinant Nbs were first pre-incubated with the trypsin resin for approximately 2-5 mins to remove the N terminal T7 tag, which is highly reactive to the crosslinker. Nb was incubated with RBD in the pH 7.4 buffer at 4°C for 1 hr to allow the formation of the complex. The reconstituted complexes were then cross-linked with 2 mM disuccinimidyl suberate (DSS, ThermoFisher Scientific) for 25 min at 25°C with gentle agitation. The reaction was then quenched with 50 mM ammonium bicarbonate (ABC) for 10 min at room temperature. After protein reduction and alkylation, the cross-linked samples were separated by a 4–12% SDS-PAGE gel (NuPAGE, Thermo Fisher). The regions corresponding to the monomeric, cross-linked species (~45-50 kDa) were sliced and digested in-gel with trypsin and Lys-C {Shi, 2014 #5336; Shi, 2015 #5385; Xiang, 2020 #5722}. After efficient proteolysis, the cross-link peptide mixtures were desalted and analyzed with a nano-LC 1200 (Thermo Fisher) coupled to a Q Exactive™ HF-X Hybrid Quadrupole-Orbitrap™ mass spectrometer (Thermo Fisher). The cross-linked peptides were loaded onto a Picochip column (C18, 3 μm particle size, 300 Å pore size, 50 μm × 10.5 cm; New Objective) and eluted using a 60 min LC gradient: 5% B–10% B, 0 – 2 min; 10% B – 40% B, 2 – 50 min; 40% B–100% B, 50 – 60 min; mobile phase A consisted of 0.1% formic acid (FA), and mobile phase B consisted of 0.1% FA in 80% acetonitrile. The QE HF-X instrument was operated in the data-dependent mode. The top 4 most abundant ions (with the mass range of 350 to 2,000 and the charge state of +4 to +7) were fragmented by high-energy collisional dissociation (normalized HCD energy 30). The target resolution was 120,000 for MS and 15,000 for MS/MS analyses. The quadrupole isolation window was 1.6 Th and the maximum injection time for MS/MS was set at 300 ms. After the MS analysis, the data was searched by pLink for the identification of cross-linked peptides. The mass accuracy was specified as 10 and 20 p.p.m. for MS and MS/MS, respectively. Other search parameters included cysteine carbamidomethylation as a fixed modification and methionine oxidation as a variable modification. A maximum of three trypsin missed-cleavage sites was allowed. Initial search results were obtained using the default 5% false discovery rate, estimated using a target-decoy search strategy. The crosslink spectra were manually checked as previously described {Xiang, 2020 #69;Shi, 2015 #70;Shi, 2014 #71;Kim, 2018 #72}.

### Integrative structural modeling

Structural models for Nbs were obtained using a multi-template comparative modeling protocol of MODELLER {Sali, 1993 #50}. Next, we refined the CDR3 loop {Fiser, 2003 #51} and selected the top 5 scoring loop conformations for the downstream docking in addition to 5 models from comparative modeling. Each Nb model was then docked to the RBD structure (PDB 6LZG) by an antibody-antigen docking protocol of PatchDock software that focuses the search to the CDRs and optimizes CXMS-based distance restraints satisfaction {Schneidman-Duhovny, 2012 #52; Schneidman-Duhovny, 2020 #53}. A restraint was considered satisfied if the Ca-Ca distance between two DSS cross-linked residues is within 28Å. The models were then re-scored by a potential statistical SOAP {Dong, 2013 #54}. The antigen interface residues (distance <6Å from Nb atoms) among the top 10 scoring models, according to the SOAP score, were used to determine the epitopes. The precision was estimated based on the convergence among the ten top-scoring models which was measured as the average RMSD between all the pairs.

### Crystallization, data collection, and structure determination of RBD-Nb20 complex

Crystallization trials were performed with the Crystal Gryphon robot (Art Robbins). The RBD-Nb20 complex was crystallized using the sitting-drop vapor diffusion method at 17 °C. The crystals were obtained in conditions containing 100 mM sodium cacodylate pH 6.5 and 1 M sodium citrate. For data collection, the crystals were transferred to the reservoir solution supplemented with 20% glycerol before freezing in liquid nitrogen. X-ray diffraction data were collected at the Advanced Photon Source (APS) beamline 23IDB of GM/CA with a 10μm-diameter microbeam. The data were processed using HKL2000 {Otwinowski, 1997 #59}. Diffraction data from six crystals were merged to obtain a complete dataset with a resolution of 3.3 Å.

The structure was determined by the molecular replacement method in Phaser {McCoy, 2007 #60} using the crystal structures of RBD (PDB 6LZG) and an Nb (V_H_H-72, PDB 6WAQ) as search models. The initial model was refined in Phenix {Adams, 2010 #61}and adjusted in COOT {Emsley, 2004 #62}. The model quality was checked by MolProbity {Williams, 2018 #63}. The final refinement statistics were listed in Table S3.

Nb21 comparative modeling was done using the Nb20 structure as a template in MODELLER. All structure visualization figures were prepared using UCSF ChimeraX {Goddard, 2018 #55}.

### Thermostability analysis of Nbs

Nb thermostability was measured by differential scanning fluorimetry (DSF). To prepare DSF samples, Nbs were mixed with SYPRO orange dye (Invitrogen) in PBS to reach a final concentration of 12 μM. The samples were analyzed in triplicate using a 7900HT Fast Real-Time PCR System (Applied Biosystems) as previously described {Shen, 2020 #31}. The melting point was calculated by the first derivatives method {Niesen, 2007 #56}.

### Nb stability test

Nbs were dissolved in 20 mM HEPES, 150 mM NaCl, pH 7.5 at 0.4 mg/ml. 50% of the proteins were lyophilized by snap-freezing in liquid nitrogen, and speed-vac dried. ddH2O was used to reconstitute the proteins. The other 50% were aerosolized using a portable mesh atomizer nebulizer (MayLuck). The aerosolized droplets were collected in a microcentrifuge tube. SEC and pseudovirus neutralization assays were performed as described above to assess the properties and activities of the Nbs.

### Data availability

The coordinates and structure factors for SARS-CoV-2 RBD with Nb20 have been deposited in the Protein Data Bank under the accession codes PDB 7JVB.

